# Connectomic features underlying diverse synaptic connection strengths and subcellular computation

**DOI:** 10.1101/2021.08.19.456845

**Authors:** Tony X. Liu, Pasha A. Davoudian, Kristyn M. Lizbinski, James M. Jeanne

## Abstract

Connectomes generated from electron microscopy images of neural tissue unveil the complex morphology of every neuron and the locations of every synapse interconnecting them. These wiring diagrams may also enable inference of synaptic and neuronal biophysics, such as the functional weights of synaptic connections, but this requires integration with physiological data to properly parameterize. Working with a stereotyped olfactory network in the *Drosophila* brain, we make direct comparisons of the anatomy and physiology of diverse neurons and synapses with subcellular and subthreshold resolution. We find that synapse density and location jointly predict the amplitude of the somatic postsynaptic potential evoked by a single presynaptic spike. Biophysical models fit to data predict that electrical compartmentalization allows axon and dendrite arbors to balance independent and interacting computations. These findings begin to fill the gap between connectivity maps and activity maps, which should enable new hypotheses about how network structure constrains network function.

## INTRODUCTION

The brain coordinates perception and action using networks of diverse neurons with heterogenous synaptic connections (Reid, 2012). Understanding how these networks operate requires a synthesis of structural and functional information (Bargmann, 2012; Briggman and Bock, 2012; Denk et al., 2012; Lee and Reid, 2011). Theoretical network models can unify diverse sources of information into a common framework, but to make the most informed computational predictions, modeled neurons and synapses should correspond to real neurons, and synapses and should obey known biophysical principles (Friedrich and Wanner, 2021; Litwin-Kumar and Turaga, 2019; Marder, 1998). With densely reconstructed connectivity maps from electron microscopy (EM) images, real network anatomy can now constrain the structure of large network models, improving biological accuracy (e.g., Eschbach et al., 2020; Tschopp et al., 2018; Vishwanathan et al., 2021; Wanner and Friedrich, 2020; Zarin et al., 2019). However, other sources of biophysical diversity, such as variations in synaptic connection weights (i.e., the physiological strength between synaptic partners) or subcellular compartmentalization are not fully constrained by connectivity maps. Instead, these are often treated as free parameters to be optimized or assumed to be homogeneous. These unconstrained variables make connectivity maps difficult to link directly to activity maps, because different biophysical parameters can produce divergent network function (Bargmann and Marder, 2013).

It has been proposed that a small number of anatomical features measurable from a complete connectome might be sufficient to predict true synaptic connection weights and to determine when axons or dendrites need to be explicitly modeled (Litwin-Kumar and Turaga, 2019). Examining this requires a direct comparison of physiology and anatomy across a diverse set of neurons and synapses. Compellingly, postsynaptic density size linearly relates to unitary excitatory postsynaptic potential (uEPSP) amplitude for a single type of neocortical synapse in mouse (Holler et al., 2021), but scaling this to more diverse populations is limited, in part, by the current lack of a full wiring diagram of the mouse brain. However, a recent connectome details the diversity of neural morphologies and synaptic connections throughout half of the central brain of the adult fruit fly, *Drosophila melanogaster* (the “hemibrain”; Scheffer et al., 2020).

Here, we ask how anatomical patterns of connectivity and neuron morphology predict physiological connection strengths and subcellular electrical compartmentalization in the *Drosophila* brain. We focus on the lateral horn, an anatomically diverse olfactory network (analogous to the mammalian cortical amygdala) mediating innate responses to odors (Dolan et al., 2019; Fişek and Wilson, 2014; Frechter et al., 2019; Kohl et al., 2013; Ruta et al., 2010). In this network, second-order projection neurons (PNs) connect to third-order lateral horn neurons (LHNs; Amin and Lin, 2019). Because connectivity and morphology are stereotyped (Bates et al., 2020b; Schlegel et al., 2021), we can study the anatomy and physiology of the same neurons and connections in different flies. We thus leveraged the hemibrain connectome along with a recent dataset of electrical recordings of identified PN-LHN uEPSPs (Jeanne et al., 2018), to make comparisons between anatomy and physiology at the elemental level of single neurons and single synaptic connections.

We report three main findings. First, physiological connection weight (i.e., uEPSP amplitude) increases linearly with synapse density (the number of synapses per unit area of postsynaptic membrane) across a diverse sample of connection types. Second, connection weight (measured at the soma) decreases with synaptic distance along the “inter-arbor cable” that connects the axon and dendrite arbors. In contrast, synaptic distance within each arbor has little impact. Subcellular electrical compartmentalization can thus be predicted by a neuron’s anatomical arrangement of arbors and cables. Third, biophysical models constrained by anatomical and physiological data predict that LHNs are shaped to maximize input gains in each arbor, while still allowing for passive interactions between arbors. This highlights a potential functional role for the inter-arbor cable in regulating subcellular computation. Together, these findings reveal important links between connectivity maps and activity maps, which should enable construction of more biologically naturalistic network models.

## RESULTS

### PN-LHN connections and LHN sizes reflect anatomical diversity of the whole brain

Within the lateral horn, PN axons carrying information from each of ∼50 olfactory antennal lobe glomeruli converge and diverge onto hundreds of morphologically distinct LHN types (one example pair in Figure 1A), providing an anatomical foundation for diverse transformations of the incoming PN odor code (Bates et al., 2020b; Dolan et al., 2019; Fişek and Wilson, 2014; Frechter et al., 2019; Jeanne et al., 2018; Jeanne and Wilson, 2015; Schlegel et al., 2021). In *Drosophila*, connections between neurons typically consist of multiple synaptic contacts, each of fairly similar physical size (Scheffer et al., 2020; Tobin et al., 2017; we use “synapse” to refer to individual contact sites, and “connection” to refer to all the synapses between two neurons). We focus on uniglomerular cholinergic (excitatory) PNs (subsequently referred to simply as “PNs”), which were the PN types studied physiologically (Jeanne et al., 2018).

**Figure 1.**
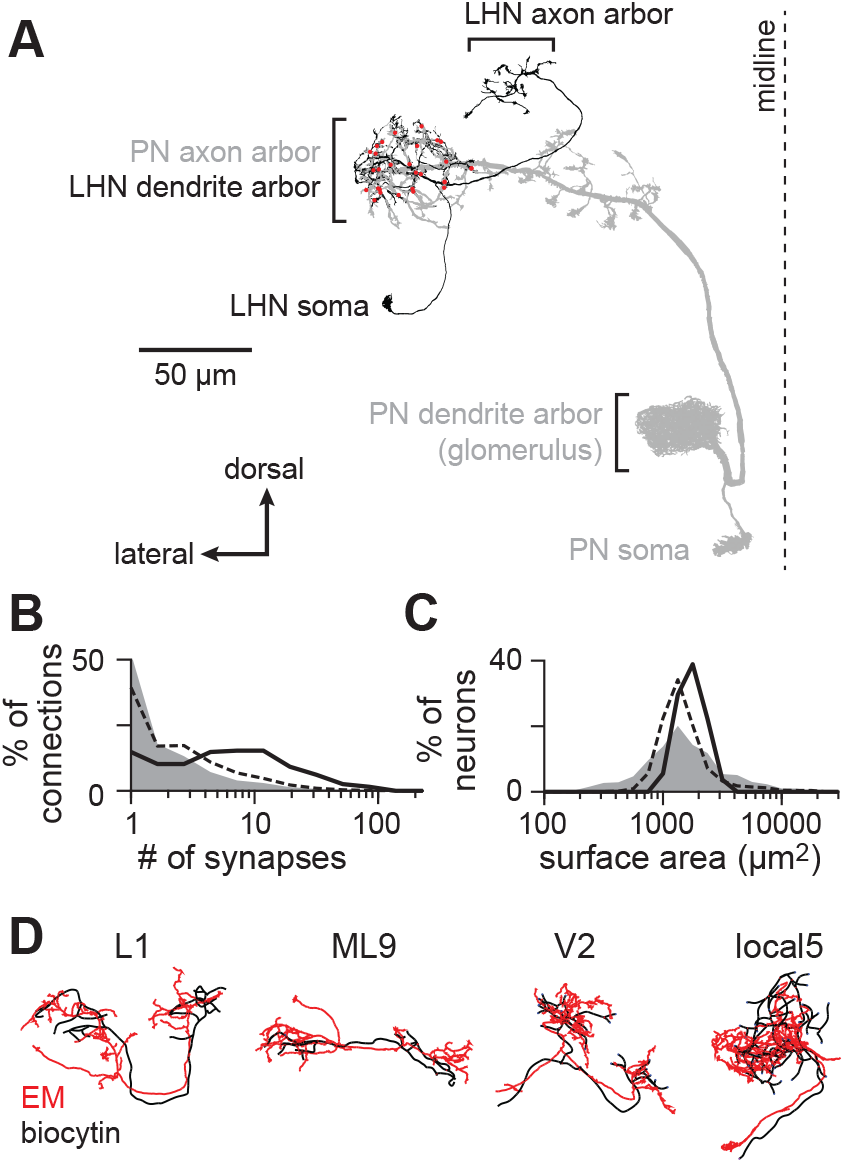
Matching diverse LHNs between light and electron microscopy datasets. **(A)** Morphology of a single PN-LHN connection. The PN’s axon arbor targets the lateral horn, where it forms multiple synapses (red points) onto an LHN’s dendrite arbor. **(B)** Distribution of the number of synapses per connection for a representative sample of connections across the hemib-rain (random sample of 214,245 connections, gray), for all 17,506 PN-LHN connections (dashed line), and for the 146 PN-LHN connections matched to physiology data (solid line). **(C)** Distribution of the total membrane surface area for a representative sample of neurons across the hemibrain (random sample of 1000 neurons, gray), for all 1496 LHNs (dashed line), and for the 54 LHNs matched to physiology data (black line). **(D)** Example matching morphologies. Black: biocytin fill (physiology dataset). Red: EM morphology (anatomy dataset).

The hemibrain connectome (Scheffer et al., 2020) annotates 100 PNs and 1496 LHNs, with 17,506 connections between them (most of the 149,600 PN-LHN pairs are not connected). The distribution of synapse counts per connection spans more than two orders of magnitude (Figure 1B). The distribution of LHN surface areas spans about one order of magnitude (Figure 1C). Distributions of both PN-LHN synapse counts and LHN surface areas are similar to distributions for connections and neurons throughout the brain (Figure 1B,C). The diversity of neurons and connections in the lateral horn is therefore representative of the fly brain as a whole.

### Cellular morphology identifies PNs and LHNs between datasets

To study anatomy and physiology of the same neurons, we used the hemibrain connectome in conjunction with previous current clamp recordings of LHNs during stimulation of PNs (Jeanne et al., 2018). Taking advantage of the stereotyped anatomy of the antennal lobe and lateral horn (Bates et al., 2020b; Schlegel et al., 2021), we identified the same PN and LHN types across these two datasets. PN types were matched by the glomerulus innervated by their dendrites, which uniquely and unambiguously identifies them (Marin et al., 2002; Wong et al., 2002). Glomeruli vary in the number of PNs innervating them, but we only considered glomeruli with one PN, which allowed us to resolve single uEPSPs in LHN recordings (Methods). LHN types were matched by quantitatively comparing biocytin fills with neuron morphologies in the hemibrain (Methods; Figure 1D, S1; Tables S1 and S2; Costa et al., 2016). In total, we matched 12 PNs and 13 LHN types between datasets.

The anatomical properties of this subset were representative of the entire lateral horn. Comparing connections from this subset of PN-LHN pairs to those from all pairs in the lateral horn shows a similarly wide distribution of synapse counts (Figure 1B) and the surface areas of these LHNs are similar to surface areas of all LHNs (Figure 1C). In addition, PN-LHN connections of the same type have similar synapse counts (Figure S2A) and LHNs of the same type have similar surface areas (Figure S2B).

### Unitary PN-LHN synaptic potentials are physiologically diverse

We analyzed LHN membrane voltages measured in response to two-photon optogenetic activation of single PNs (Figure 2A; Jeanne et al., 2018). Activation of individual PNs produces distinct uEPSP waveforms in the LHN (each corresponding to one PN spike; Figure 2B). We extracted each uEPSP for each PN-LHN connection within a recording (Methods; Figure 2C). Overall, uEPSPs were measurable and reliably detected over multiple LHN recordings for 29 unique PN-LHN pairs. No synaptic responses were evident in 106 pairs, so we considered these to have uEPSP amplitudes of 0 mV. The remaining pairs had inconsistent or unquantifiable synaptic responses and were omitted (Methods), yielding a population of 135 unique PN-LHN pairs matched to anatomy and with reliable estimates of physiology.

**Figure 2.**
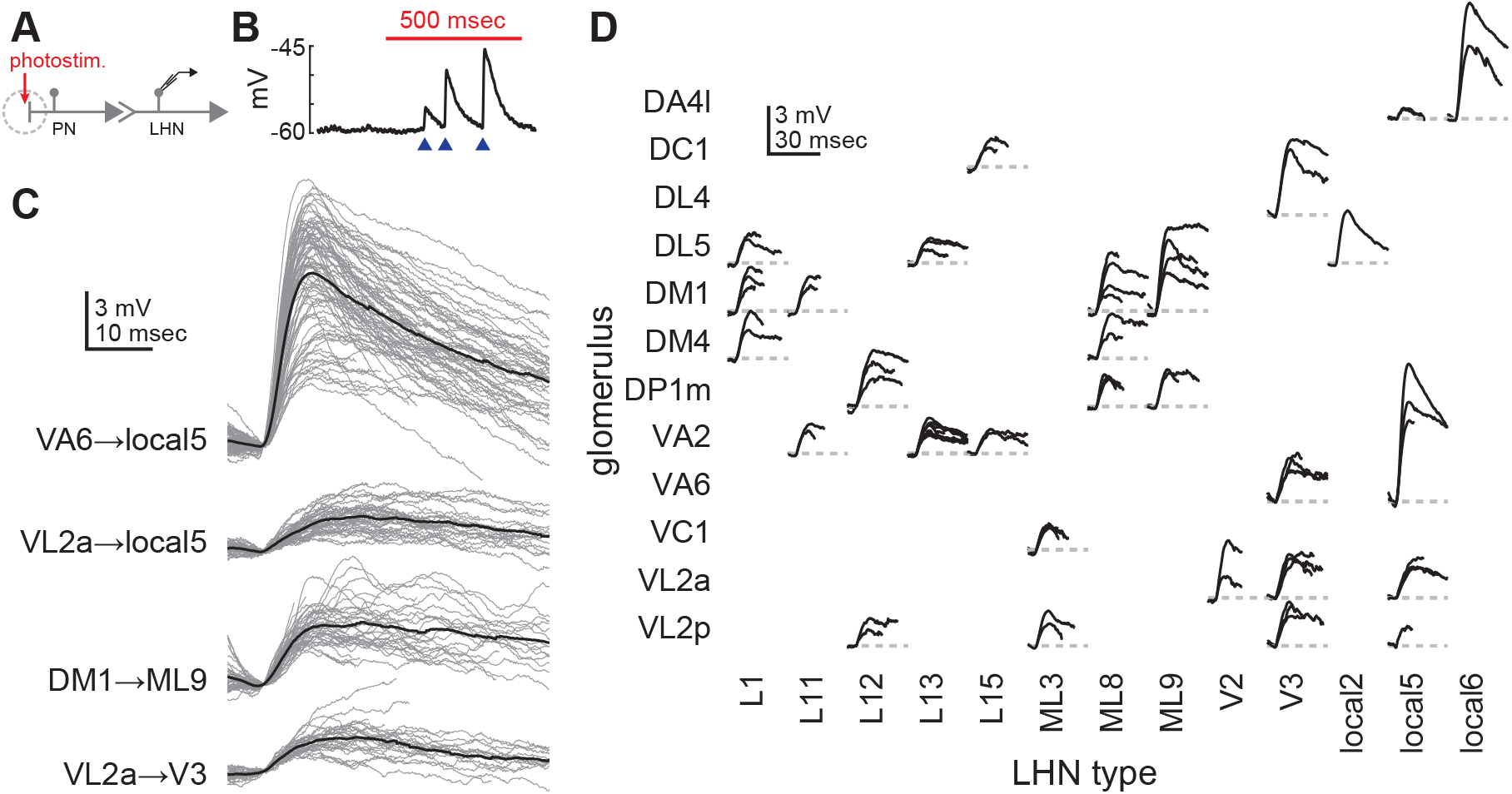
PN-LHN connections have diverse physiological weights. **(A)** Schematic of experimental configuration. PN dendrites are photostimulated during patch-clamp recording from single LHNs. **(B)** Example trace recorded from an LHN during PN stimulation (red bar). Blue arrowheads denote single uEPSPs corresponding to single PN spikes. **(C)** Overlaid traces (gray) of all individual uEPSPs (aligned by start time) detected from individual recordings of four different PN-LHN connections. Each black trace is the mean of all uEPSPs from one recording of one connection type. **(D)** Mean uEPSP traces for all identified connections. Multiple recordings (from different flies) of the same connection type are overlaid. Variability between mean uEPSP amplitudes for different connection types is greater than for repeated recordings within the same types (ANOVA, F = 8.8, p = 3.3 × 10^−13^).

Mean uEPSP amplitudes spanned a ∼10-fold range (0.4mV to 6.6mV), with distinct and characteristic values for different connection types, but with some variability (Figure 2B-D). Variability of individual uEPSP waveforms within a single recording (e.g., Figure 2C) likely arises from physiological factors that modify synaptic function, such as short-term plasticity or stochastic transmitter release. Occasional variability of mean uEPSP waveforms for the same PN-LHN connection type across recordings may reflect fly-to-fly variability in synapse counts, or differences in long-term plasticity or neuromodulatory state. Despite this variability, clear differences in waveforms between PN-LHN connection types were still observable (Figure 2D). We thus averaged mean uEPSPs across recordings for each PN-LHN connection type to compare to anatomical data from the same connections.

### Anatomical diversity linearly predicts physiological diversity of synaptic connections

Next, we compared the anatomical and physiological strengths of PN-LHN connections. In the quiescent (*ex vivo*) condition of our recordings, LHNs likely operate in a passive regime. Thus, synaptic potentials should be proportional to the product of synaptic conductance, driving force, and membrane resistance (Johnston and Wu, 1995). Moreover, because our measured uEPSPs are relatively small compared to the synaptic reversal potential (−10 mV; Methods), driving force should be approximately constant. Therefore, synapse density (synapse count normalized by LHN surface area) should predict uEPSP amplitude, because surface area is inversely proportional to membrane resistance (assuming uniform membrane conductance). Consistently, synapse density predicted average uEPSP amplitude well (Figure 3A, solid line; r^2^ = 0.77, p = 1.0 × 10^−43^). Importantly, this correlation persisted even without the largest connections or the pairs with both 0 synapses and 0 mV uEPSP amplitudes (Figure S3) and fit with data for connections in the antennal lobe and mushroom body (Figure S4). Thus, anatomical features can predict average physiological weights for diverse connection types (without invoking active properties), in a way that may be conserved throughout the fly brain.

**Figure 3.**
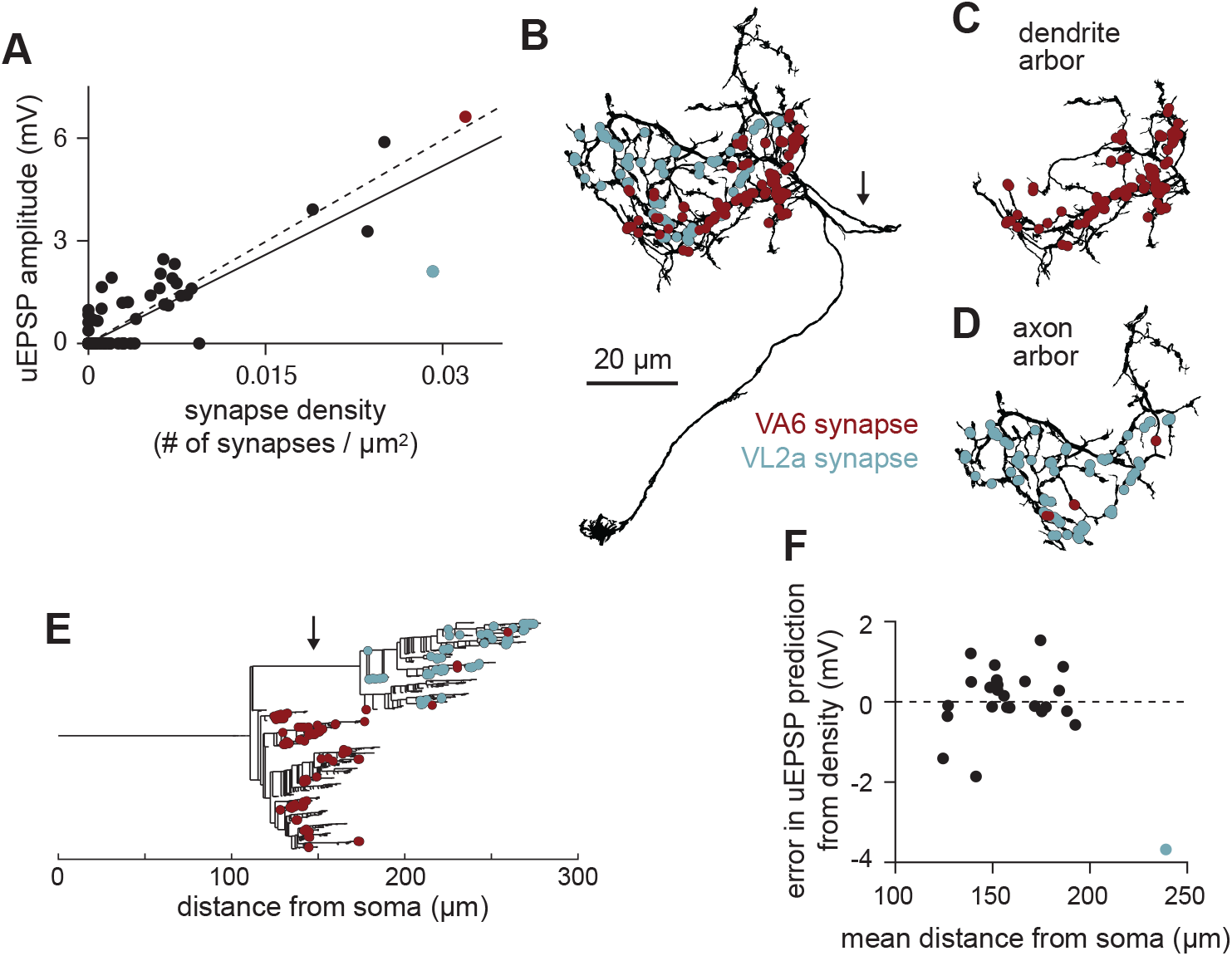
Anatomical diversity predicts physiological diversity of PN-LHN connections. **(A)** Scatter plot of uEPSP amplitude vs. synapse density for all connections in our sample. Red and blue points correspond to the VA6 and VL2a connections onto local5. Note that many points are overlaid at the origin, corresponding to connections with no synapses detected in the hemibrain and no uEPSPs detected in physiology. Solid line is linear fit including the axo-axonic VL2a-local5 connection; dashed line excludes this connection. **(B)** Morphology of a local5 neuron (bodyId 5813105722), with all VA6 and VL2a synapse locations labeled. Arrow identifies the inter-arbor cable, which follows a meandering path to connect the two intermingled arbors. **(C)** As in (B) but showing just the dendrite arbor. Almost all the VA6 synapses are formed onto this part of the neuron. **(D)** As in (B) but showing just the axon arbor. All of the VL2a synapses are formed onto this part of the neuron. **(E)** Dendrogram showing the branching structure of the same local5 neuron as in (D)-(F). Almost all of the synapses onto the axon arbor are more distant from the soma than the synapses onto the dendrite arbor, which is due primarily to the inter-arbor cable (identified by arrow). **(F)** Residual error in uEPSP amplitude from synapse density prediction, showing no linear correlation with mean distance of synapses from the soma. uEPSP amplitudes of dendritic inputs are thus not strongly dependent on distance.

Interestingly, one outlier stood out in this analysis: the connection from the VL2a PN onto local5 LHNs. This uEPSP was substantially smaller than predicted by its high synapse density (blue point in Figure 3A). Interestingly, local5 LHNs also received a connection from the VA6 PN, which had a similarly high synapse density, but a large uEPSP amplitude (red point in Figure 3A). We also noticed that the VL2a-local5 uEPSP had relatively slow kinetics (Figure 2C). The time to the peak amplitude for the VL2a connection was longer (11.9 ± 1.7 msec) than for the corresponding VA6 connection (7.1 ± 0.2 msec). However, the uEPSPs between VL2a and a different LHN (“V2”) exhibited a short time to peak (7.9 ± 0.3 msec), indicating that slow kinetics cannot be due to slower synaptic release from the VL2a PN.

The co-occurrence of small amplitudes and slow kinetics in the VL2a-local5 connection suggests unusually strong cable filtering, which attenuates and delays synaptic potentials (Rall, 1967). We thus asked whether the VL2a PN targets local5 more distally than the VA6 PN. The neurites of local5, like those of most local LHNs, organize into two polarized arbors: one biased towards input synapses (dendrite) and the other biased towards output synapses (axon). The arbors of local5 extensively intertwine with each other (Figure 3B), but viewed in isolation, a segregated pattern of synapses emerges. VA6 synapses almost exclusively target the dendrite arbor (Figure 3C), while VL2a synapses exclusively target the axon arbor (which is more distant from the soma; Figure 3D,E). Thus, the longer distance to the soma for the VL2a connection predicts greater cable filtering and reduced somatic uEPSP amplitude, when compared to the VA6 connection.

Given this role of synaptic distance, we reasoned that it might also impact uEPSP amplitudes for other PN-LHN connections. Surprisingly, after regressing out the effect of synapse density from uEPSP predictions, the residual variability showed no obvious relation to mean synapse distance, other than for the VL2a-local5 connection (Figure 3F). Because all other connections in our dataset target LHN dendrites, this points to some aspect of arbor identity (axon vs. dendrite), instead of distance alone, as an essential determinant of postsynaptic filtering. In light of this, we recomputed the prediction of average uEPSP amplitude from synapse density using only the dendritic connections. This yielded a stronger correlation (r^2^ = 0.84; Figure 3A, dashed line). Collectively, these results show that synapse density and location both matter for determining somatic uEPSP amplitudes, but that the role of location is not a simple linear function of distance.

### Passive compartmental models accurately predict diverse uEPSP amplitudes

To better understand how synapse location and LHN morphology impact uEPSP amplitudes, we turned to biophysical modeling. First, we built single-compartment models for every LHN in our sample by jointly optimizing membrane resistance and capacitance per unit area, and conductance per synapse to fit uEPSP amplitudes (Methods). These parameters were constrained to be uniform across LHNs, with whole-cell properties varying solely due to variability in neuron size. As expected, these models forced a compromise between underpredicting dendritic and overpredicting axonal uEPSPs (RMS error = 0.95mV; Figure 4A,B). This illustrates the insufficiency of single compartment models for neurons with multiple arbors.

**Figure 4.**
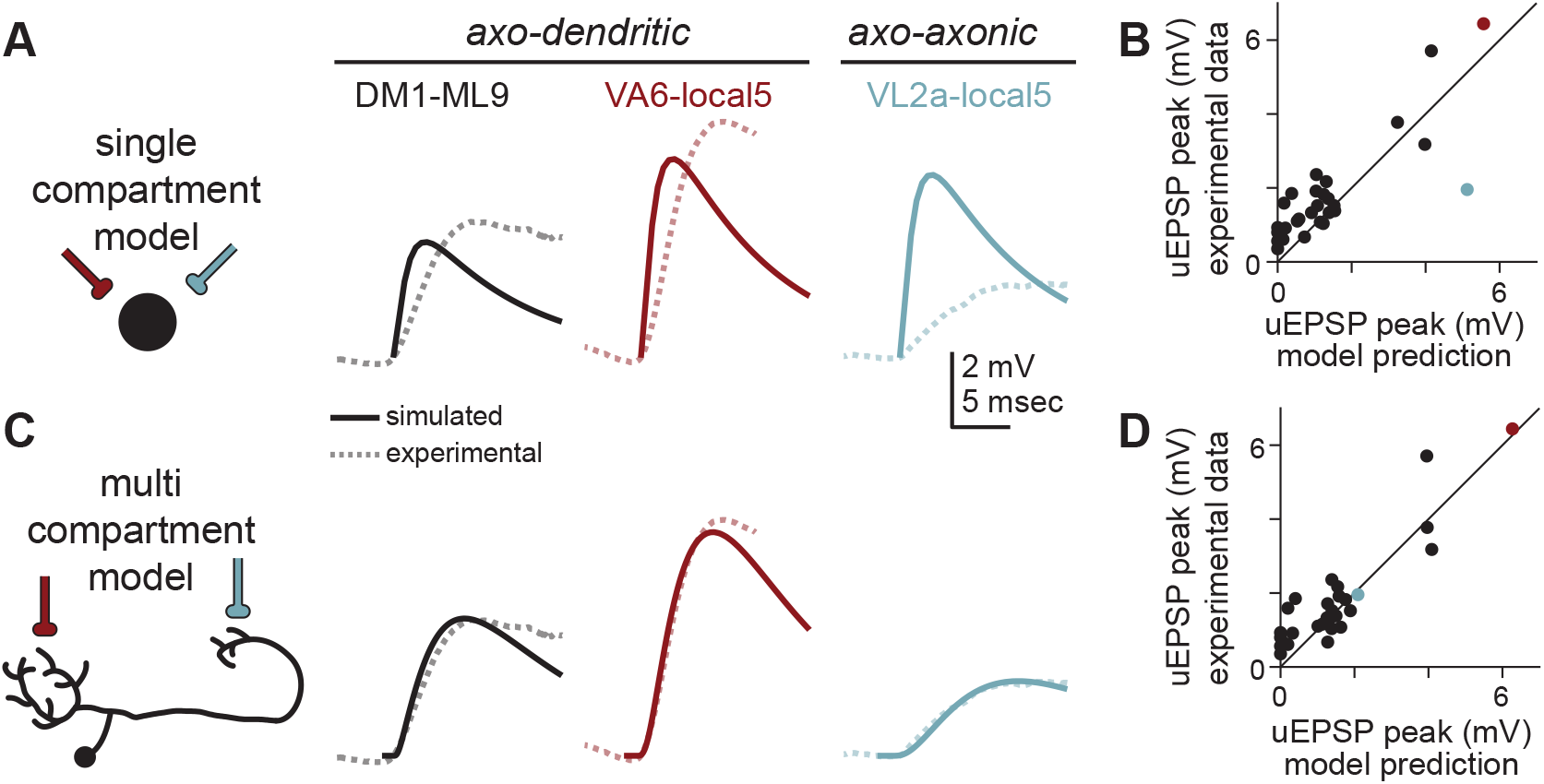
Multi-compartment models accurately predict uEPSP amplitudes better than single compartment models. **(A)** A single compartmental (point) model (schematized at left) fit to all PN-LHN uEPSPs in our sample (dashed lines) produces somatic uEPSP waveforms (solid lines) that underpredict true uEPSP amplitudes for axo-dendritic connections and overpredicts the axo-axonic connection. **(B)** Peak somatic uEPSP amplitudes for all connections in our sample (with detectable uEPSPs) compared to predictions from the single compartment models. Note the blue point (corresponding to the VL2a-local5 connection) is well below the unity line, while most other points (including small amplitude connections) are above the line. **(C)** As in (A) but for multi-compartment models. The optimized model fits axo-dendritic connections as well as the axo-axonic connection. **(D)** As in (B) but for the multi-compartment model.

We then fit multi-compartment models (with morphology and synapse locations explicitly constrained by hemibrain data) by additionally optimizing axial resistance (Methods). Again, all parameters were uniform across LHNs, so models only vary due to morphology and synapse location. These models predicted the amplitude of uEPSPs across our dataset with better accuracy (RMS error = 0.68mV; Figure 4C,D). This indicates that differences in uEPSP amplitude for connections onto different arbors can be parsimoniously explained by differences in passive filtering arising from neuronal morphology, requiring neither active conductances nor cell-type or subcellular specificity of passive properties.

### Models predict that arbors democratize synaptic efficacies, while inter-arbor cables stratify them

We next used multi-compartment models to understand the complex relation between anatomical synapse locations and physiological synapse amplitude, focusing exclusively on local LHNs because they receive PN input onto both dendrite and axon arbors. Using the parameter values determined by the fits to data described above, we built models for 49 local LHNs, including local5 (Methods). These models allowed us to “stimulate” single synapses and “record” from any neuronal location. Accordingly, we decomposed uEPSPs into the constituent potentials evoked by individual synapses, called miniature EPSPs (mEPSPs), measured either at the synaptic location or at the spike initiation zone (SIZ; located near the base of the dendritic arbor in fly neurons; Gouwens and Wilson, 2009; Figure 5A). We refer to the amplitude of the mEPSP at the SIZ as “synaptic efficacy,” and note that voltage at the soma largely tracks voltage at the SIZ, but with additional passive filtering.

**Figure 5.**
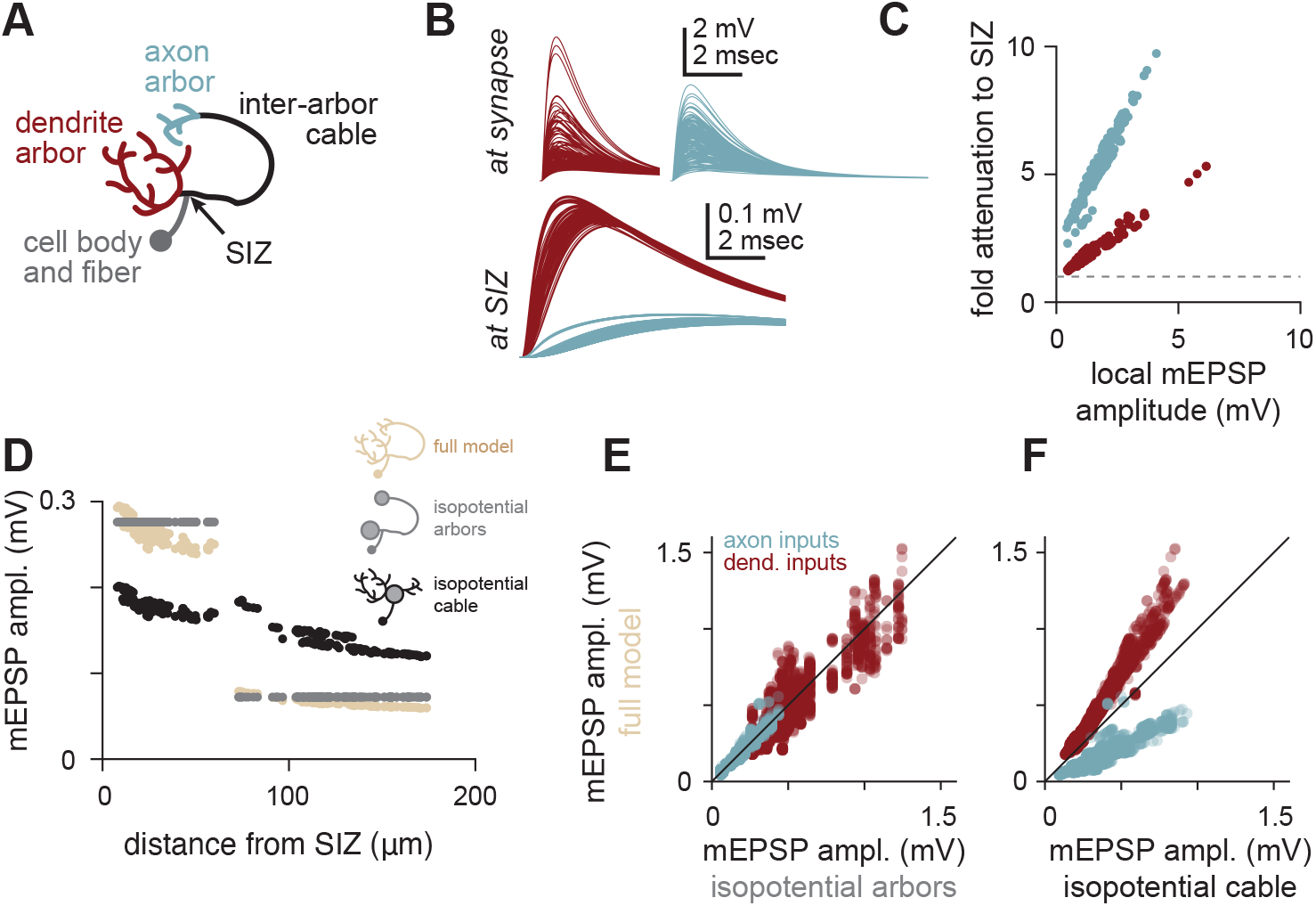
Arbors democratize and cables stratify synaptic efficacy. **(A)** Schematic of arbors, cables, and spike initiation zone (SIZ) in local LHNs. **(B)** Top: traces of simulated mEPSPs measured at each synapse (red, dendritic; blue, axonal) onto a local5 LHN (same example as Figure 3). Local mEPSPs are large and similar between arbors. Bottom: traces of simulated mEPSPs measured at the SIZ. Differences in mEPSPs are apparent between arbors. **(C)** Local mEPSP amplitudes relate linearly to their voltage attenuation en route to the SIZ. Within each arbor, synapses with larger local depolarizations also face proportionally larger attenuation. This proportionality is different for each arbor. Red points are dendritic mEPSPs, blue points are axonal mEPSPs. **(D)** mEPSP amplitudes for all ePN synapses onto the same local5 LHN, as a function of distance from the SIZ (tan). The inter-arbor cable introduces a discontinuity in this relationship: while the most proximal axonal synapses are nearly the same distance from the SIZ as the most distal dendritic synapses (∼60 μm), they evoke strikingly smaller mEPSP amplitudes. Grey points are the same measurements, but for a model with isopotential arbors. Black points are for a model with an isopotential inter-arbor cable. **(E)** mEPSP amplitudes simulated with the full model vs. the isopotential arbor model for all synapses across all local LHNs in our sample. The isopotential arbor model was a good predictor of the mEPSPs in the full model (r^2^ = 0.87). **(F)** mEPSP amplitudes simulated with the full model vs. the isopotential cable model for all synapses across all local LHNs in our sample. The isopotential cable model introduces systematic biases in mEPSP amplitudes, making it a less good predictor (r^2^ = 0.47).

We first recorded mEPSPs at synaptic sites, to ask whether local morphological differences yield systematic electrical differences between axon and dendrite arbors. In the local5 LHN model, mEPSPs at synaptic sites were diverse, with no apparent differences between arbors (Figure 5B). Because the synaptic conductance is the same for all synapses in our model, all diversity in mEPSPs must come from variations in local input resistance. This means that local electrical properties at each synapse are variable, but that axons and dendrites have similar amounts of variation.

We then recorded the same mEPSPs at the SIZ. There we saw the opposite pattern: mEPSPs were uniform for synapses on the same arbor, but with clear differences for synapses on different arbors (Figure 5B). This suggests subcellular processing by LHNs democratizes synaptic efficacies within arbors, but stratifies efficacies (i.e. introduces characteristic differences) between arbors.

To better understand the within-arbor democracy, we compared local mEPSP amplitudes with the fold attenuation to the SIZ (Figure 5C). This revealed strong linear correlations for all synapses within each arbor (but with different slopes and intercepts for different arbors). Therefore, synapses within an arbor generate widely different local depolarizations, but these differences are mostly normalized by compensatory voltage attenuation *en route* to the SIZ (tan points in Figure 5D). Synaptic efficacies from the “perfect” democracy implemented by a model with isopotential arbors were minimally different from the full model (gray points in Figure 5D), which was consistent for synapses across all local LHNs we studied (Figure 5E, Methods). This democratization of synaptic inputs (without active conductances) likely arises from the tapering of neurites and the relatively shallow and wide tree structure of LHN arbors, which have also been observed in other cell types (Cuntz et al., 2007; Jaffe and Carnevale, 1999; Roth and Hausser, 2001).

In contrast to the democratizing effect of the arbors, the inter-arbor cable stratifies synaptic efficacies. This is evident from the fact that the isopotential arbor model (devoid of all morphological detail within each arbor) retains the differences between arbors (Figure 5D,E). Notably, in a model with an isopotential cable, all synaptic efficacies took on intermediate values (and varied as a smooth function of distance; Figure 5D,F). Therefore, the long and thin shape of the inter-arbor cable both reduces the synaptic efficacy of axonal synapses, and increases the efficacy of dendritic synapses, by introducing substantial electrotonic compartmentalization between arbors. We also note that parallel effects occur for mEPSP latency (Figure S5). Stratifying synaptic efficacies between arbors but not within arbors results from cable theory (Rall, 1962). The inter-arbor cable is flanked by current sinks (i.e., arbors) on either end, so it acts like an infinite cable (where voltage attenuates exponentially with distance). In contrast, the shorter cables that make up each arbor have a large current sink only on one end and are sealed on the other. Thus, arbors act more like semi-infinite cables with closed boundaries, meaning that nearly all synaptic current flows towards the arbor root (with minimal voltage attenuation).

### PN connections onto local LHNs mostly target individual arbors

The most obvious way for subcellular compartmentalization of LHNs to participate in transforming the incoming PN odor code is if individual PNs from different glomeruli target different arbors. We have shown already that VA6 and VL2a PNs target different arbors of local5 LHNs, but it is not clear whether this patterning occurs frequently. We therefore asked whether PN axons selectively target individual arbors across our larger sample of local LHNs (Figure 6A). Strikingly, 72% of connections (of 5 synapses or larger) targeted one arbor nearly exclusively (≥90% of synapses targeting one arbor), much more than expected from random targeting (Figure 6B).

**Figure 6.**
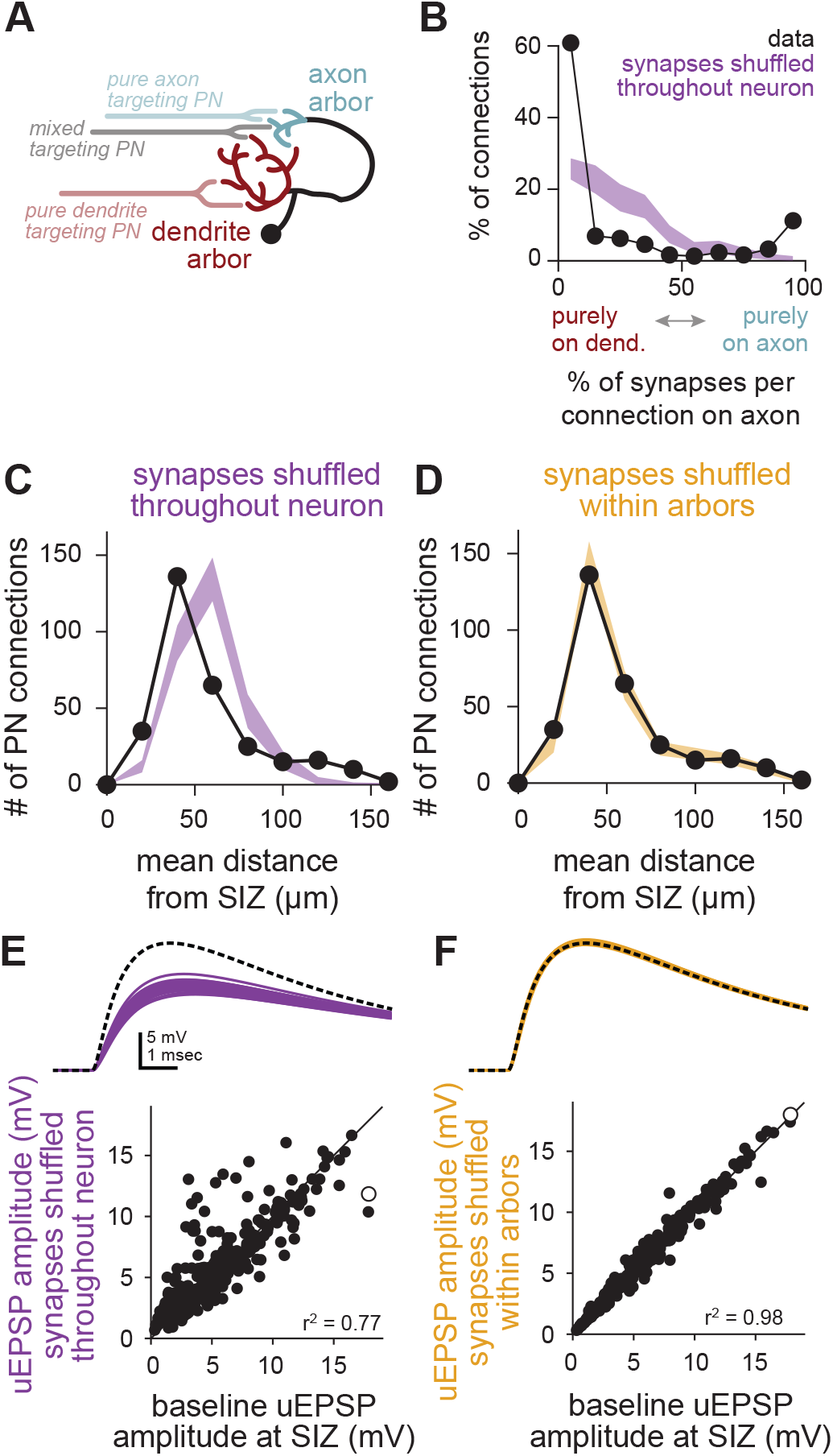
PN connections target synapses to specific arbors but lack sub-arbor organization. **(A)** Schematic of PNs targeting axons and dendrites of local LHNs. **(B)** Most PN connections (with ≥5 synapses) purely target one local LHN arbor or the other. Compared to a null distribution generated by shuffling all PN synapse labels throughout each neuron (purple band denotes the 95% confidence interval), connections are strongly biased towards targeting a single arbor (black points). Most dendrites receive more synapses than axons, so the null distribution skews left. **(C)** Arbor-specific targeting of PN connections broadens the distribution of mean distances to the SIZ across connections onto 49 local LHNs. Black points are the true distribution of mean distances for each connection. Purple band denotes the 95% confidence interval for randomly shuffled synapse locations throughout the whole neuron. **(D)** As in (C) but compared to a null model of synapse locations shuffled within arbors (orange band: 95% confidence interval). **(E-F)** Shuffling synapse locations throughout the neuron (E) perturbs uEPSP amplitudes more than shuffling only within arbors (F). Example uEPSP traces at top (dashed black line denotes trace for real synapse locations; each colored line denotes one shuffle) correspond to open circles in scatter plots below.

We next asked whether connections exhibit sub-arbor specificity by comparing average distances to the SIZ for all synapses making up each PN to local LHN connection. Mean distances for these connections spanned a broad range, but this range was underpredicted by a null model where PN synapse locations were permuted throughout each LHN (Figure 6C). In contrast, the distribution of distances was well-matched by a null model where synapse locations were permuted only within arbors (Figure 6D). This illustrates that PN synapses precisely target specific arbors, but less precisely target specific locations within arbors.

Because PNs may also target their synapses to sub-arbor structures that are uncorrelated with distance (e.g., individual branches), we quantified the impact of shuffling synapse locations on the simulated uEPSP amplitude for each connection. When synapse locations were permuted throughout the entire neuron (i.e., across both arbors), the resulting uEPSP amplitudes were less predictive of the true amplitudes (r^2^ = 0.77; Figure 6E) than when synapses were only permuted within arbors (r^2^ = 0.98; Figure 6F). This further supports the conclusion that synapses from PNs closely track the compartmentalization of LHNs, largely isolating their connections to either the dendrite or axon arbor, but with minimal organization within each arbor.

### Models predict that inter-arbor cables establish independent, robust, and maximal local input gain

Thus far, we have considered how synaptic inputs depolarize the SIZ. While this region is of clear importance, each arbor may also perform local functions. For instance, direct excitation of axons may modulate spike-evoked neurotransmitter release or even drive graded release without spikes (Graubard et al., 1980).

However, local computation requires some degree of electrotonic independence between arbors, which appears at odds with our observation of inter-arbor voltage propagation. How do LHNs strike a balance between independence and interaction between arbors?

For arbors to maintain some independence, synaptic currents should mostly charge their local membranes, without much leak to distant membranes. To see if this is the case, we used our compartmental models to investigate each arbor’s local input gain. For the same synaptic conductance, an arbor with higher input gain will exhibit a larger local depolarization. Input gain is thus well approximated in our models by the mean mEPSP amplitude (which minimally impacts driving force) across all synapses. We measured amplitudes at each arbor’s primary branch point, to ensure we capture arbor-specific democratization and to focus on differences between arbors. We found that the input gain was larger in axon arbors than in dendrite arbors (Figure 7A), inversely tracking differences in arbor size of local LHNs (Figure 7B). Accordingly, local arbor membrane resistance almost completely predicted the variability in local input gain for both axon and dendrite inputs (r^2^ = 0.87; Figure 7C). Thus, each arbor operates with its own independent input gain, with relatively little synaptic current is lost to distal membranes.

**Figure 7.**
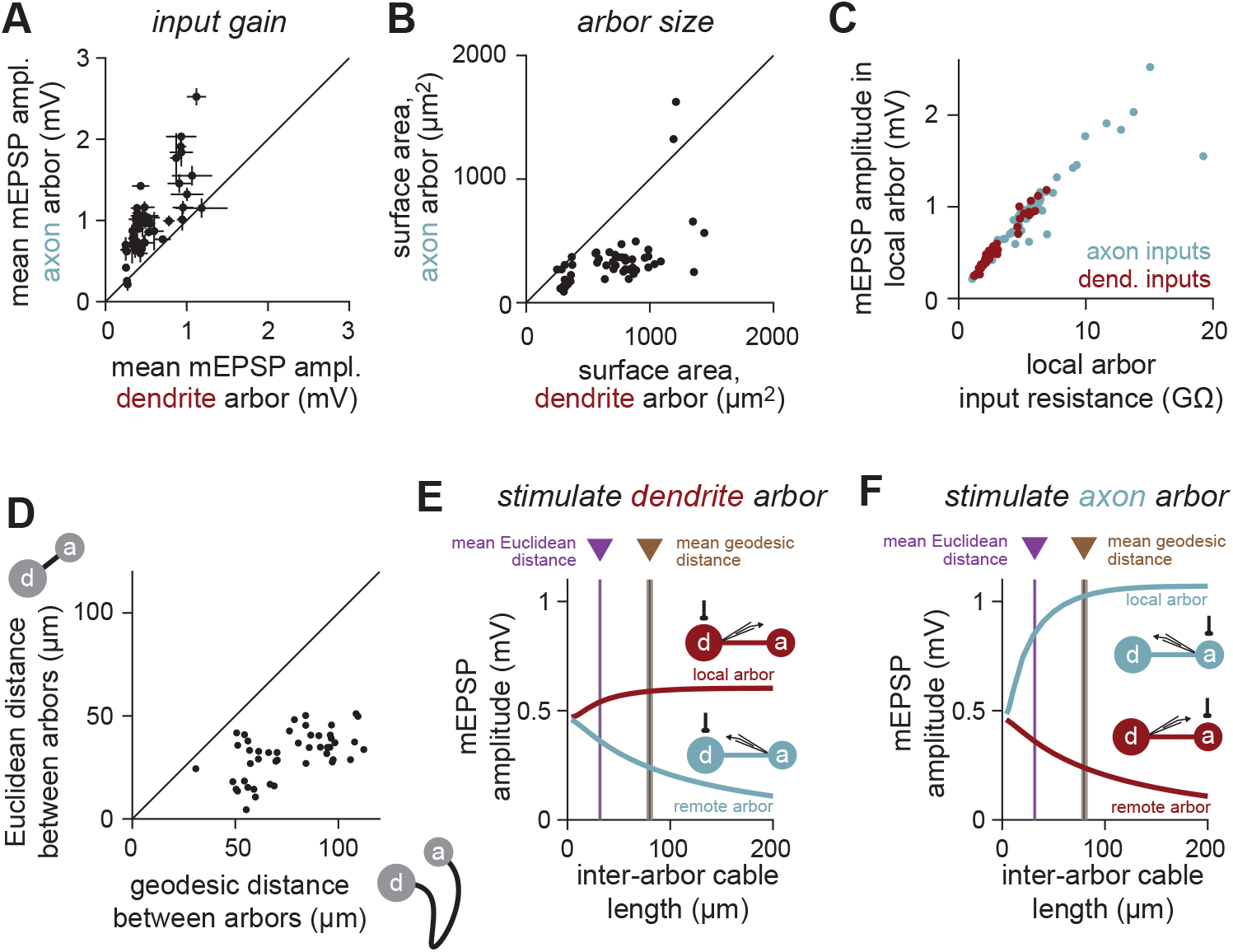
Inter-arbor cable lengths balance independence of and interaction between arbors. **(A)** The local arbor input gain (measured as mEPSP amplitude at the first branch point of each arbor) is larger in axons than dendrites (i.e., the same size synaptic conductance depolarizes the axon more than the dendrite). Each point denotes one local LHN (error bars are S.D.). **(B)** The surface area of axon arbors is smaller than dendrite arbors. **(C)** Input gain (local mEPSP amplitude) can be predicted by the local arbor input resistance, for both axon and dendrite inputs. Consequently, each arbor is minimally affected by distant current sinks, such as the other arbor. The shared slope comes from the shared specific membrane resistance. **(D)** Geodesic distance (path length) of local LHN inter-arbor cables are longer than the Euclidean distance (straight line path) between arbors. **(E-F)** Input gain (local arbor mEPSP amplitude) in the barbell (simplified isopotential arbor) model of an average local LHN depends strongly on the length of the inter-arbor cable. The true (geodesic) length of the cable balances maximizing local input gain while minimizing signal loss between arbors.

Membrane resistances, however, can fluctuate dramatically due to synaptic activity or intrinsic plasticity (Destexhe et al., 2003; Zhang and Linden, 2003). We therefore wondered whether local input gain is robust to changes in remote membrane resistance. We simulated an average local LHN with a simplified “barbell” model: a single compartment for each arbor and a multi-compartment cable connecting them (Methods).

Utilizing the barbell model, we systematically varied the membrane resistance of the remote arbor while measuring local mEPSP amplitude. Across a wide range of remote membrane resistances, local mEPSP amplitudes were unchanged (Figure S6), indicating that local input gain is robust to variations in remote arbor membrane resistance.

We next looked more closely at inter-arbor cables. These are especially interesting in local LHNs (e.g., arrow in Figure 3B), because they meander along paths that deviate considerably from the shortest (Euclidean) path between arbors (2.8±1.7 times longer; Figure 7D). This hints that cable length may be tuned to perform computational functions – such as balancing independent input gains against inter-arbor interactions – rather than simply to connect inputs to outputs in different spatial locations.

To test this idea, we manipulated the inter-arbor cable length in the barbell model. Lengthening the cable increased local input gain (top curves in Figure 7E,F) by impairing the other arbor’s ability to siphon off current. However, input gain saturated once the cable was long enough to resemble a semi-infinite cable (Rall, 1959). Interestingly, the true (geodesic) mean cable length matched the point of saturation, such that further lengthening would have vanishing effects on local input gain. Lengthening the inter-arbor cable also decreased mEPSP amplitudes measured from the remote arbor. However, these continued to decrease after the local mEPSPs saturated (bottom curves in Figure 7E,F).

Collectively these simulations suggest that the independence of each arbor is established by an inter-arbor cable that is just long enough to maximize local input gains. However, the cable is also short enough to permit passive exchange of information, allowing axonal input to contribute to depolarizing the SIZ. Thus, the inter-arbor cable balances independence and interaction between arbors, potentially expanding the computational repertoire of single neurons.

### Predicted functions of compartmentalized neurons

How might the independent input gains in each arbor affect how a neuron transforms inputs into outputs? While our model did not allow for direct simulation of active properties, we used prior measurements of spike threshold and general properties of transmitter release to predict how input to each arbor controls output. Revisiting the barbell model of the average local LHN, we made functional predictions for three configurations of activation: input only to the dendrite arbor, input only to the axon arbor, and input to both arbors together.

First, when inputs arrive only onto the dendrite arbor, the LHN operates in a “feedforward spiking drive” mode. With a spike threshold 15mV above rest at the SIZ (Fişek and Wilson, 2014; Jeanne and Wilson, 2015), strong dendritic input (0.03 synapses/μm^2^) can rapidly evoke spikes, while a weaker input (0.0075 synapses/μm^2^) does not (Figure 8A). Varying synapse density and input spike rate shows that only the strongest connections evoke spikes on their own; weaker connections would require coincident activity of multiple inputs (Figure 8B).

**Figure 8.**
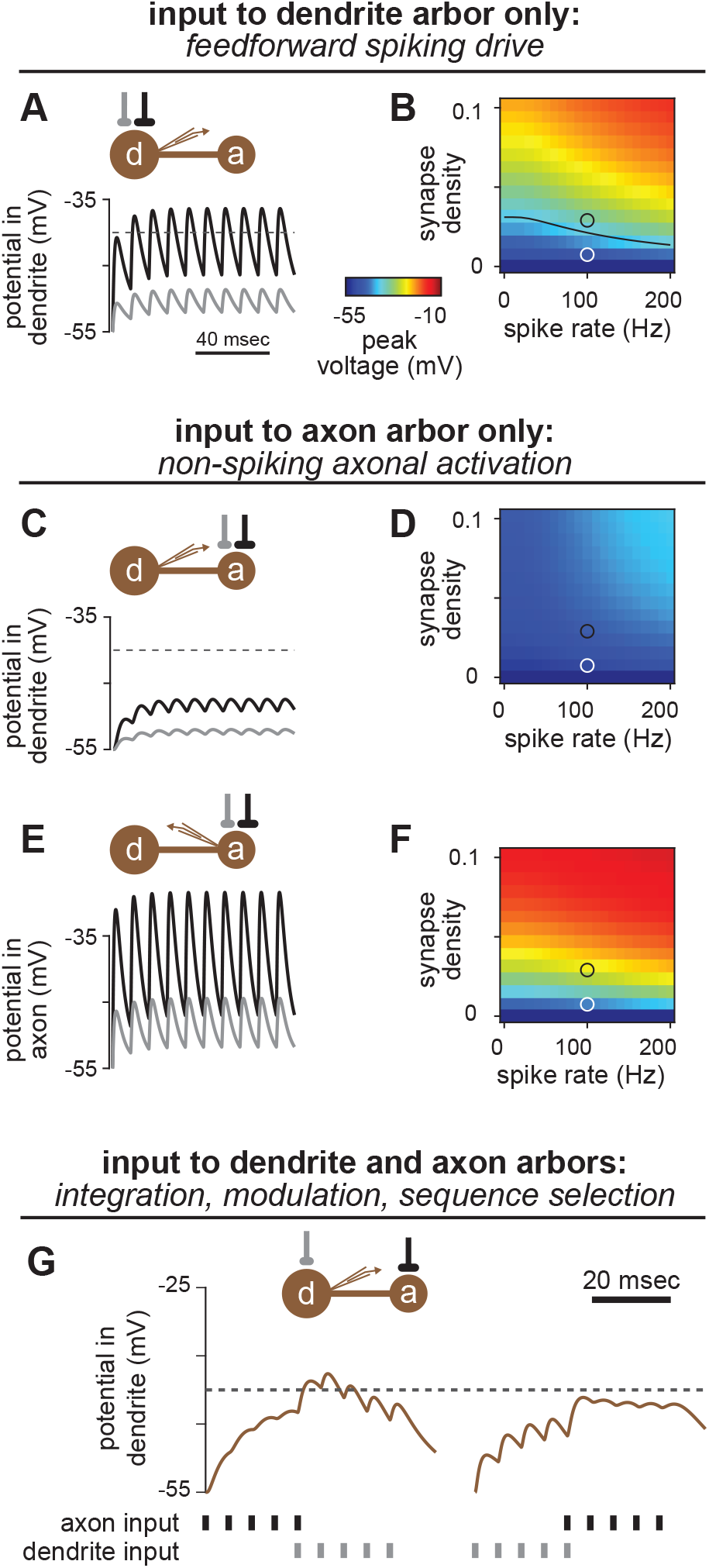
Predicted functions of compartmentalized neurons. **(A-B)** Input to the dendrite arbor alone should readily drive spikes in local LHNs. (A) In the average local LHN (simulated with the barbell model; Methods), a relatively strong connection (0.03 synapses/μm2, black trace) driven at 100Hz depolarizes the SIZ above spike threshold (−40mV). A weaker connection (0.0075 synapses/μm2, gray trace) does not reach spike threshold. (B) Peak voltages obtained for dendritic stimulation for a range of synapse densities and input spike rates. Black line in the plot corresponds to the spike threshold of -40mV. The black and gray circles correspond to the black and gray traces in (A), respectively. **(C-D)** Input to the axon arbor alone cannot drive spikes. (C) Axonal connections with the same synapse densities and spike rate as in (A) generate much smaller depolarizations at the SIZ. Colors are as in (A). (D) None of the simulated synapse densities or spike rates could depolarize the dendrite to spike threshold. **(E-F)** Input to the axon arbor alone substantially depolarizes the axon, potentially driving graded release.. Plots are as in (C-D), but with recording in the axon arbor. **(G)** Temporal delays along the inter-arbor cable create selective responses to different input sequences. A small spike packet impinging on the axon that precedes a spike packet impinging on the dendrite elicits a stronger response (left curve) in the dendrite than the opposite sequence (right curve). The connection onto the axon is stronger than the connection onto the dendrite, to compensate for attenuation.

Second, when inputs arrive only onto the axon arbor, the LHN operates in a “nonspiking axonal activation” mode. Even the strongest physiologically plausible inputs (simultaneous activation of all PN inputs to an LHN axon) with high spike rates are insufficient to depolarize the SIZ to spike threshold (Figure 8C,D). However, these inputs strongly depolarize the axon directly (Figure 8E,F), potentially causing graded (spike-independent) transmitter release, which starts at voltages of ∼10-15mV above rest in invertebrates (Angstadt and Calabrese, 1991; Burrows and Siegler, 1978; Graubard, 1978; Graubard et al., 1980).

Third, when inputs arrive on both the dendrite and axon, the LHN operates in one of several modes. In the simplest case, these inputs may simply integrate. In another case, direct depolarization of the axon might alter the amount of neurotransmitter released by action potentials, reminiscent of cortical neurons (Alle and Geiger, 2006; Shu et al., 2006). Finally, the delay introduced by propagation along the inter-arbor cable (Figure S5) may enable sequence-selective responses. With spike generation occurring near the dendrite arbor, synaptic input sequences that activate the axon before the dendrite cause larger depolarizations of the SIZ than the opposite sequences (Figure 8G). Together, these simulations provide a framework for generating explicit hypotheses about how neuronal morphology shapes subcellular computation within the context of network-level function.

## DISCUSSION

### Predicting synaptic function from connectomic data

Our results show that much of the diversity in mean physiological connection weights between PNs and LHNs can be explained by anatomical properties measurable in EM images. The number of synapses and LHN surface area successfully predict somatic uEPSP amplitudes for dendritic connections, but severely underpredicts amplitudes for axonal connections. However, a more complex model incorporating neural morphology and synapse locations accurately predicted the physiology of both connection types. This highlights the insufficiency of neural point models (which ignore morphology) for predicting synaptic and neural function.

It is perhaps surprising that such a strong correspondence occurs with purely passive models, given the plethora of voltage-gated ion channels expressed in neurons (Littleton and Ganetzky, 2000). However, the quiescent network state (*ex vivo* preparation) and the minimal stimulation (single spike resolution in single neurons) are both favorable conditions for remaining in a passive regime. We also note a slight tendency for underprediction of some of the smaller uEPSP amplitudes (Figure 3A, 4D), which might indicate some active boosting at the low end, or some sublinear integration at the high end of connection weights. Interestingly, a similar phenomenon has been observed in triplet recordings of PNs and LHNs (Fişek and Wilson, 2014). Nevertheless, our passive models should serve as a foundation in which to investigate the role of active properties.

We found it striking that the relationship between dendritic synapse density and uEPSP amplitude was largely linear, and that connections in the antennal lobe and mushroom body were consistent with this relationship. This suggests a conservation of fundamental biophysics across neurons and synapses. Notably, the capacitance of neural membrane and the resistance of intracellular medium are fairly consistent in different neurons and species (Borst and Haag, 1996). Although membrane resistance is more variable due to differences in ion channel expression and activation, the quiescent network state likely quenches many of these differences. The baseline synaptic conductance is likely fairly uniform across synapses in the *Drosophila* brain (Scheffer et al., 2020; Tobin et al., 2017), but does vary in a use-dependent manner (Hige et al., 2015; Kazama and Wilson, 2008). We thus anticipate that our predictions of synaptic function from PN-LHN connections will apply to other connections in the fly brain, but that some features, such as synaptic plasticity, will remain idiosyncratic.

Our results also highlight the importance of comparing structure and function with single neuron and single spike resolution. While we found that connectomic data accurately predicted uEPSPs evoked by single spikes in single neurons, a recent study (using the same data sources) reported lower predictive power for compound EPSP amplitudes evoked by multiple spikes in multiple neurons (Schlegel et al., 2021). Future efforts to link connectivity and physiology may thus face challenges if sufficient physiological resolution is not obtained, but our results suggest that biophysical models could fill the gap in certain instances. For example, spatial and temporal integration across neural populations can be explicitly modeled to match commonly used experimental measures, such as voltage imaging from neuropil containing multiple cells or cell types.

### Arbors and cables in cellular computation

The morphologies of neurons are famously diverse (Ramon y Cajal, 1995). Yet while morphology affects synaptic efficacy in some neurons (Stuart et al., 2016), it may play a minimal role in others (Otopalik et al., 2017). For LHNs, we find evidence of both phenomena: fine morphology within arbors has limited impact on synaptic efficacy, while the inter-arbor cable has a major impact.

More specifically, our results show that individual arbors passively integrate synaptic input democratically. This occurs because large local variations in mEPSP amplitude are mostly offset by compensatory variations in voltage attenuation. The electrotonic structure of central *Drosophila* neurons may thus be similar to the dendrites of cerebellar Purkinje cells, which orchestrate a similar dendritic democracy with passive mechanisms in a heavily branched arbor (Roth and Hausser, 2001). Interestingly, this configuration has been attributed to the lack of a central trunk neurite (Jaffe and Carnevale, 1999; Figure S7), so the branching characteristics of fly neuron arbors may be a mechanism to achieve uniform synaptic efficacy without special spatial patterning of ion channel expression or synaptic conductances (Cuntz et al., 2008). In addition, because most PN-LHN connections target a single arbor with multiple spatially distributed synapses, much of the residual variability due to synapse location will average out for larger connections. Single arbors may therefore be fundamental “units” of computation in *Drosophila* neurons, which can spatially intermingle even within the same brain region.

Inter-arbor cables strike a balance between interaction and independence between arbors. Interaction enables neurons to compare inputs arriving on different arbors. This is especially relevant, because axon and dendrite arbors receive their own complements of synaptic inputs. LHNs are thus reminiscent of coincidence-detector neurons in the auditory brainstem, where input from each ear impinges on different dendritic arbors, allowing the connecting cable to compare timing (Agmon-Snir et al., 1998). Inter-arbor cables in local LHNs are longer than necessary to connect the arbors, enabling discrimination of temporal sequences on behaviourally relevant scales of ∼10 msec (Egea-Weiss et al., 2018; Sehdev et al., 2019). In contrast, independence between arbors can enable some functions to remain arbor-specific. For instance, arbor-specific structural plasticity (Liu et al., 2017) or active conductances (e.g., voltage-gated potassium channels; Diao et al., 2010; Rogero et al., 1997) could implement different transformations within each arbor prior to comparison via the inter-arbor cable. Such a configuration could enable more complex computations such as multiplication (Gabbiani et al., 2002).

The abstraction of intricate morphologies into arbors and cables should prove useful for studying other neurons. Even within the fly brain, a wide range of configurations exist, including neurons with one arbor and no cable (some local neurons; Chou et al., 2010; Schlegel et al., 2021), neurons with one arbor and one cable (Kenyon cells without axonal branching), three arbors with interposed cables (optic lobe neurons; Fischbach and Dittrich, 1989; Yang et al., 2016), and 2-dimensional arrays of dozens of arbors (amacrine neurons; Meier and Borst, 2019). An intriguing possibility is that the arbor and cable configuration largely determines the passive biophysics of these neurons, which could provide a simple organizing framework for predicting the function of neurons with diverse neuron morphologies.

### From connectivity maps to activity maps

The pairing of network connectivity maps with knowledge of neuronal and synaptic physiology provides a foundation to formulate hypotheses about activity maps, because assumptions about the function of each component can be calibrated (Litwin-Kumar and Turaga, 2019). We take important steps in this direction by showing how synapse densities predict uEPSP amplitudes and how morphology predicts subcellular computation. Moreover, we demonstrate that a simplified compartmental model (the barbell model) can balance biological accuracy with computational tractability. Incorporating these results into simulations of large neural networks should allow the formulation of more precise mechanistic hypotheses about the function of previously unexplored brain circuits.

An important goal for the future will be to incorporate additional sources of knowledge to constrain other properties, such as synaptic plasticity, active conductances, and neuromodulation. For example, short-term plasticity can correlate with the number of presynaptic vesicles or the location of a synapse along the dendrite (Abrahamsson et al., 2012; Xu-Friedman and Regehr, 2004). A systematic comparison of synaptic ultrastructure with synaptic plasticity may therefore reveal other structural patterns that predict function (e.g., Eckstein et al., 2020). Active properties of neurons are less likely to be predictable from ultrastructure but could be predicted from proteomics and transcriptomics (Croset et al., 2018; Davie et al., 2018). Relating physiological measurements to these data across cell types could be used to calibrate estimates of active biophysical properties. Neuromodulation can reconfigure entire networks, yielding different functions in different conditions (Harris-Warrick and Marder, 1991). In the fly, dopamine alters physiological synaptic strengths, but it is not clear if such changes are visible in EM images (Cohn et al., 2015; Hige et al., 2015). High resolution mapping of cellular and synaptic biochemical processes would thus be a valuable companion to a connectome (Aso and Rubin, 2020).

Another goal will be to incorporate ongoing refinement to wiring diagrams into model calibrations. This is important, because some regions of the hemibrain connectome still have incompletely traced connections (Scheffer et al., 2020). In addition, this connectome also lacks information on electrical synapses, glia, and some subcellular structures, which have important functional roles. As this information becomes available, the calibration of cellular and synaptic predictors can be continually adjusted and improved.

While connectivity maps of increasingly large brain volumes bring new opportunities for understanding network organization, predicting function from structure remains famously difficult (Marder, 2015). While we have focused here on the function of individual components (i.e., neurons and synapses), a central challenge will be identifying how the operation of entire networks depends on those components. Although physiological properties of synapses and neurons are most easily characterized in quiescent network states (e.g., *ex vivo*), many network operations occur in highly active states (e.g., *in vivo*). (Destexhe et al., 2003). The combination of connectivity maps with validated models of synaptic and neuronal function should help to bridge this gap, by generating testable predictions of how the anatomy and physiology of neurons and synapses constrain activity maps during behavior.

## ACKNOWLEDGEMENTS

We thank Stephen Plaza for help with the hemibrain dataset and Greg Jefferis for assistance with neuroanatomical analysis. We thank Michael Higley, Damon Clark, Mehmet Fişek, and members of the Jeanne Lab for helpful discussions and for comments on the manuscript. This work was supported by NIH grants R01 DC018570 and R01 NS116584, the Richard and Susan Smith Family Award for Excellence in Biomedical Research, the Klingenstein-Simons Fellowship Award in Neuroscience, and an innovative research grant from the Kavli Institute for Neuroscience at Yale University to J.M.J. T.X.L. was supported by the Yale College Dean’s Research Fellowship and the Rosenfeld Science Scholars Program. P.A.D was supported by NIH medical scientist training grant T32GM136651. K.M.L. was supported by a James Hudson Brown – Alexander Brown Coxe Postdoctoral Fellowship in the Medical Sciences at Yale University School of Medicine and NIH fellowship F32 DC019521.

## AUTHOR CONTRIBUTIONS

J.M.J. conceived and designed the study with input from all authors. K.M.L. and J.M.J. analyzed electrophysiology data. P.A.D. analyzed connectomic data. T.X.L. designed and implemented the compartmental models. J.M.J. designed and implemented the barbell models. T.X.L., P.A.D., K.M.L., and J.M.J. wrote the paper.

## METHODS

### Data sources and inclusion

Whole-cell patch-clamp recordings (one recording per fly) were obtained from a previously published study of physiological connectivity in the lateral horn (Jeanne et al., 2018). These data also included skeletons of each recorded neuron traced from confocal images of biocytin fills. Detailed anatomy data were obtained from the publicly accessible database of traced neurons in the “hemibrain” EM connectome (version 1.1; Scheffer et al., 2020).

To ensure reliable estimates, we only included LHN types in our analysis if at least two neurons of that type were recorded as a part of the physiology dataset (Jeanne et al., 2018). In addition, connections were excluded when they were not detected reliably (we required a connection to be detected at least two times across all recordings of that LHN type in Jeanne et al., 2018) or when distinct uEPSPs could not be identified. In two cases (DL5-local2 and VL2p-local5), uEPSPs could only be identified in one of two recordings but a connection was clearly visible (without distinct uEPSPs) in the other recording, so we included these connections using solely the uEPSPs from one recording. We also excluded from all subsequent analysis those recordings of PN-LHN connections with fewer than three uEPSP waveform samples. Of the 156 unique PN-LHN type connections, 21 were excluded for the reasons listed here, yielding 135 connections with reliable and quantifiable physiological responses.

The bilateral LHN type CML2 was omitted from analysis becuase its contralateral processes exited the hemibrain EM volume, so full morphology information was not available. LHN type ML4 was omitted because this neuron exhibited spontaneous (independent of photostimulation) synaptic activity, making it impossible to assign specific EPSPs to specific PNs. In total, the 13 included LHN types consisted of 54 individual LHN hemibrain body IDs. The bodyIds of each PN and LHN are provided in tables S1 and S2, respectively.

In the process of analyzing the electrophysiology traces, we made several minor adjustments from the connections reported in Jeanne et al. (2018). The VA7l-local6 connection reported in that study was found to be misidentified and was corrected to be from glomerulus VA7m here. Connections from DP1m to ML8 and ML9 and from VA2-L11 did not meet the time-averaged depolarization threshold requirement for detection in Jeanne et al. (2018), but nonetheless exhibited several clear uEPSPs waveforms, so we included them. Within the set of 14 LHN types, we never observed reliable and measurable connectivity from five single-PN glomeruli (VC2, VM4, DM3, VA4, and VA7l), so the PNs innervating these glomeruli were excluded from all analyses.

For analysis and modeling of the expanded population of 49 local LHNs (figure 5E,F, 6, and 7), we started with 8 of the 10 LHNs matched to our physiology data (omitting the two local2 LHNs, since they only have one arbor), and added an additional 41 neurons from the hemibrain connectome. These 41 neurons were selected because they had minimally intertwined axon and dendrite arbors (see section “manual correction of neuron morphologies” below). BodyIDs of the local LHNs used for this analysis are provided in table S3.

### Bridging biocytin light microscopy and electron microscopy data

PNs were matched between the physiology and anatomy datasets based on the glomerulus innervated, an unambiguous identifier of PN type. We focused here on only PNs of glomeruli with a singular uniglomerular cholinergic PN (see “Detection and quantification of PN-LHN uEPSPs**”** for details). In total, 12 different PNs elicited measurable uEPSPs in LHNs.

LHN types were matched between the physiology and anatomy datasets solely based on morphology by comparing biocytin fills recovered from patch-clamp recordings with traced skeletons from the hemibrain connectome. We first applied bridging transformations (Bates et al., 2020a) to bring the biocytin fills and hemibrain neuron skeletons into the same coordinate space (JRC2018F). When necessary, we mirrored biocytin fills across the midline so that all neurons appeared on the right side of the template brain (where all hemibrain LHNs reside). Due to left-right symmetry (Bates et al., 2020b), all biocytin-filled neurons included in this analysis are expected to exist on both sides of the brain. We identified all 1496 neurons in the hemibrain volume that enter the lateral horn neuropil and quantitatively compared their morphologies to the biocytin fills using NBLAST (Costa et al., 2016).

LHN cell types were defined as previously determined (Jeanne et al., 2018). Morphological matches to EM neurons were thus made without regard to cell typing that has been performed in the hemibrain connectome. This is important because those cell types were defined by both morphology and connectivity (Schlegel et al., 2021). Nonetheless, many of the matching EM neurons for each cell type closely followed hemibrain cell-type boundaries (Supplemental Table S2), although the correspondence was not one-to-one.

Owing to the inherently lower resolution of confocal microscopy relative to electron microscopy, we employed two strategies to improve comparisons of traced neurons between imaging modalities. First, we emphasized long range neurites over dendritic arbors by employing the “useAlpha” parameter of the NBLAST function. Second, we pruned the shortest processes in the EM neurons (all processes shorter than 5 μm in length).

NBLAST scores for all pairwise comparisons were computed as the mean of “forward” and “reverse” scores and were normalized to enable better comparison across diverse morphologies (Costa et al., 2016). We then selected the best matching EM neurons (top 2% based on scores) for manual inspection. We were able to readily rule-in or rule-out each of these top-scoring EM neurons as a match to the biocytin fills by directly examining similarities and differences of morphology in the common coordinate template space. Particular emphasis was placed on the course and contour of primary arbors of axonal and dendritic trees as these were consistently present in both the light and electron microscopy volumes. The corresponding EM neuron bodyIds and NBLAST scores for each cell type in the physiology dataset are provided in Table S2).

In total, 54 LHNs from the connectome could be matched to 13 LHN types defined morphologically from their biocytin fills (each LHN type consisted of neurons; Figure 1D and Table S2; mean ± s.d. NBLAST score = 0.453±0.066), corresponding to a mean NBLAST rank = 8.07 of 1496). We were thus able to identify 156 distinct PN-LHN type pairs in both the connectome and the physiology dataset.

### Manual correction of neuron morphologies

All LHN morphologies were exported directly from the hemibrain dataset via the neuprint_read_neurons function (with “heal” set to “true”) of the Natverse toolbox (Bates et al., 2020a). This produced anatomical models where each neurite was defined by many short cylindrical segments, with diameters matched to the EM data. A handful of LHNs (mostly local neurons) had heavily intertwined neurites, and the automated morphological extraction often merged or separated neurites incorrectly (as judged by visual comparison to the raw EM images using Neuroglancer). Thus, we visually inspected all LHN morphologies using NeuTube (Feng et al., 2015), and manually corrected all merge and separation errors by direct comparison to the raw EM data. Datafiles (.swc format) of all LHN morphologies included in this study (incorporating all manual corrections) are provided as Supplemental Data S1.

## QUANTIFICATION AND STATISTICAL ANALYSIS

### Detection and quantification of PN-LHN uEPSPs

Unitary excitatory postsynaptic potentials (uEPSPs) were measured from recordings of LHNs during independent photostimulation of each of 39 PN types (Jeanne et al., 2018). Stimulated PNs analyzed here were exclusively of the anterodorsal or lateral lineage and each innervate a single glomerulus. Each glomerulus is innervated by a stereotyped set of 1-8 uniglomerular cholinergic PNs (Scheffer et al., 2020; Zheng et al., 2018) and most or all PNs from each glomerulus target the same LHNs (Jeanne et al., 2018; Jeanne and Wilson, 2015; Schlegel et al., 2021). Here, we studied only the PNs of glomeruli with just one uniglomerular cholinergic PN (i.e., PN types consisting of just one neuron each). This is because glomerular photostimulation drove regularly-spaced spikes in each PN in that glomerulus, but at slightly different times and rates, evoked a complex compound EPSP in the LHN (Jeanne et al., 2018). This made reliable detection of individual distinct uEPSPs possible only for glomeruli with one PN. In these cases, regularly spaced uEPSPs were clearly discernable.

All uEPSPs were manually identified from the recorded LHN traces. The identity of the presynaptic PN was determined by the identity of the stimulated glomerulus in each trace. Each experiment included multiple stimulation sites in each glomerulus and all the trials corresponding to a given glomerulus were grouped together. Each uEPSP was identified by its characteristic asymmetric waveform (fast rise and slow decay) and its start time was manually annotated by two expert (unblinded) scorers. To minimize user-specific annotation errors, only those uEPSPs annotated by both scorers with less than 3 msec difference in start times were included (the start time was taken to be the minimum of the two times, when there was a difference).

For each recorded connection, all uEPSP waveforms were averaged to generate each trace in Figure 2D. If a connection was identified in a particular recording in Jeanne et al. (2018), but we could not reliably detect uEPSPs, that connection was excluded from computing the average, which is why a few PN-LHN pairs (DL5-local2, VL2p-local5, and DA4l-local5) have only a single trace in Figure 2D. If a connection was not identified in a particular recording in Jeanne et al. (2018), the uEPSP amplitude was considered to be zero and included in the average. To enable comparisons with anatomical measures, we averaged the peak uEPSP amplitude for all samples of the same PN and LHN types. To ensure good estimates, we only considered those connections which were detected in at least two flies (with the two exceptions noted above). Because of baseline recording noise, we could not reliably identify uEPSPs of amplitude less than 0.2mV, which may introduce a modest positive bias to the mean uEPSP amplitudes of the smallest connections. Mean uEPSP amplitudes and corresponding synapse densities are provided in Table S4.

### Compartmental modeling of LHNs

Compartmental models of LHNs were constructed in the NEURON simulation environment (Hines and Carnevale, 1997). Corrected neuron morphologies were imported into NEURON, where they were divided into short isopotential compartments using the maximal available spatial resolution from the morphological SWC file, consistent with the d-lambda rule (Carnevale and Hines, 2006). Synapse locations were determined from annotations in the hemibrain dataset and mapped to the corresponding compartment in the model using a k-nearest neighbors algorithm.

Passive biophysical properties were jointly optimized to minimize the mean squared error in uEPSP amplitude for all connections. (The peak amplitude was the most reliable metric for model fitting because the full waveform of each uEPSP was sometimes obscured by subsequent uEPSPs, due to stimulation of multiple PN spikes) A leave-one-out bootstrap procedure was implemented to validate that our parameter values were robust to variations in the set of training connections. The best fit values for specific membrane resistance (17.2 kΩ cm^2^), specific membrane capacitance (0.6 μF/cm^2^), and specific axial resistivity (350 Ω cm), were in very close agreement with experimentally constrained values for PNs and visual amacrine cells (Gouwens and Wilson, 2009; Meier and Borst, 2019; Tobin et al., 2017). As in those studies, these biophysical parameters were assumed to be uniform within and between neurons. However, whole-cell properties vary between modeled neurons due to their different sizes. The PN to LHN synaptic conductance waveform shape was modeled as a sum of two exponentials (the time constant of rise was 0.2msec and the time constant of decay was 1.1msec), matching a previous model of the ORN-PN synapse (Tobin et al., 2017). The peak amplitude of the synaptic waveform was a fourth free parameter (optimized jointly with cellular biophysical properties). The best fit value was 0.055 nS, similar to a previous fit (Tobin et al., 2017). The cholinergic synaptic reversal potential was set to -10mV (McCarthy et al., 2011). Importantly, using identical parameter values to those in prior models of *Drosophila* neurons yielded qualitatively similar findings about subcellular compartmentalization, supporting the robustness of our models (Gouwens and Wilson, 2009; Meier and Borst, 2019; Tobin et al., 2017).

Voltage measurements from specific compartments of the model were obtained by placing a virtual pipette in the appropriate location and recording in current clamp mode. Individual mEPSPs were recorded by opening up a synaptic conductance (with the waveform determined above) at the location of each individual synapse. mEPSPs were recorded either locally (in the same compartment as the synapse) or at the SIZ. uEPSPs were recorded by simultaneously opening conductances at all synapses corresponding to the presynaptic PN.

Subregions of each LHN (axon arbor, dendrite arbor, inter-arbor cable, and cell body fiber) were segmented manually using neuTube. For each arbor, the start node was identified manually, and all daughter nodes were considered part of the arbor. In some LHNs, 1-3 additional branches that were not downstream of the start node were manually identified as part of the arbor. The axon arbor was distinguished from the dendrite arbor by the longer cable connection from the cell body fiber junction, a defining characteristic of arbor identify in LHNs (Bates et al., 2020b). To verify the distinction between axon and dendrite, the ratio of input to output synapses was computed to locate the “flow centrality” of each neuron (Schneider-Mizell et al., 2016). In all cases, flow centrality agreed with our manual annotation of axon and dendrite. The inter-arbor cable was defined as the neurite spanning between the start nodes of each arbor, excluding any additional arbor branches (as defined above), and the cell body and cell body fiber. One neuron type, Local2, did not have an obvious inter-arbor cable, and had no obvious polarity of input vs. output synapse locations. We therefore determined that this neuron type only has one arbor, and no inter-arbor cable.

Perturbations to the compartmental models were carried out by adjusting the axial resistivity of specific compartments to make certain parts of the neuron isopotential. Because NEURON does not allow an axial resistivity of 0, it was set to a very small positive value (0.001 ohm cm). Simulations of neurons with isopotential subregions were carried out using Crank-Nicholson integration with fixed timestep (0.06 msec).

### Anatomical analysis of neurons and synaptic connections

Analysis of axon and dendrite targeting was conducted by splitting each neurons synaptic inputs by arbor identity. The random distribution in Figure 6B was obtained by redistributing each connection’s synapses onto axon and dendrite arbors following the distribution of all ePN synapses onto that neuron. Shuffling in Figure 6C-F was conducted by randomly permuting the synapse labels (i.e. the identity of each presynaptic partner neuron) either within each arbor, or throughout the entire neuron. This was repeated 1000 times to obtain 95% confidence intervals.

### Barbell model simulation

The barbell model is a simplified isopotential arbor model (Figure 7E,F and Figure 8) and was implemented in MATLAB. All sub-arbor shape information was abstracted away, retaining only the size of each arbor, and the length and mean diameter of the inter-arbor cable. The inter-arbor cable was simulated as a series of 1 μm long compartments with diameter matching the true mean diameter, and each arbor was simulated as a single compartment. The surface area of the cell body and cell body fiber was added to the dendrite arbor, since these structures are nearby. The SIZ was therefore isopotential with the dendrite compartment. The total surface area was nearly identical to the full model (with the only differences coming from the fixed-diameter approximation for the inter-arbor cable). In addition, all core biophysical parameters (Rm, Ri, Cm, and Gsyn) were unchanged from the best fit values obtained for the full models. Numerical integration was performed using the backward Euler method (Carnevale and Hines, 2006) with a fixed timestep of 0.01ms. The range of simulated PN spike rates used in Figure 8 (0-200Hz) approximated the known dynamic range of these neurons (Bhandawat et al., 2007).

### Statistics

Central tendencies are reported as means, and dispersions are reported as standard deviations, except for figure 6, where dispersions are reported as 95% confidence intervals.

**Figure S1:**
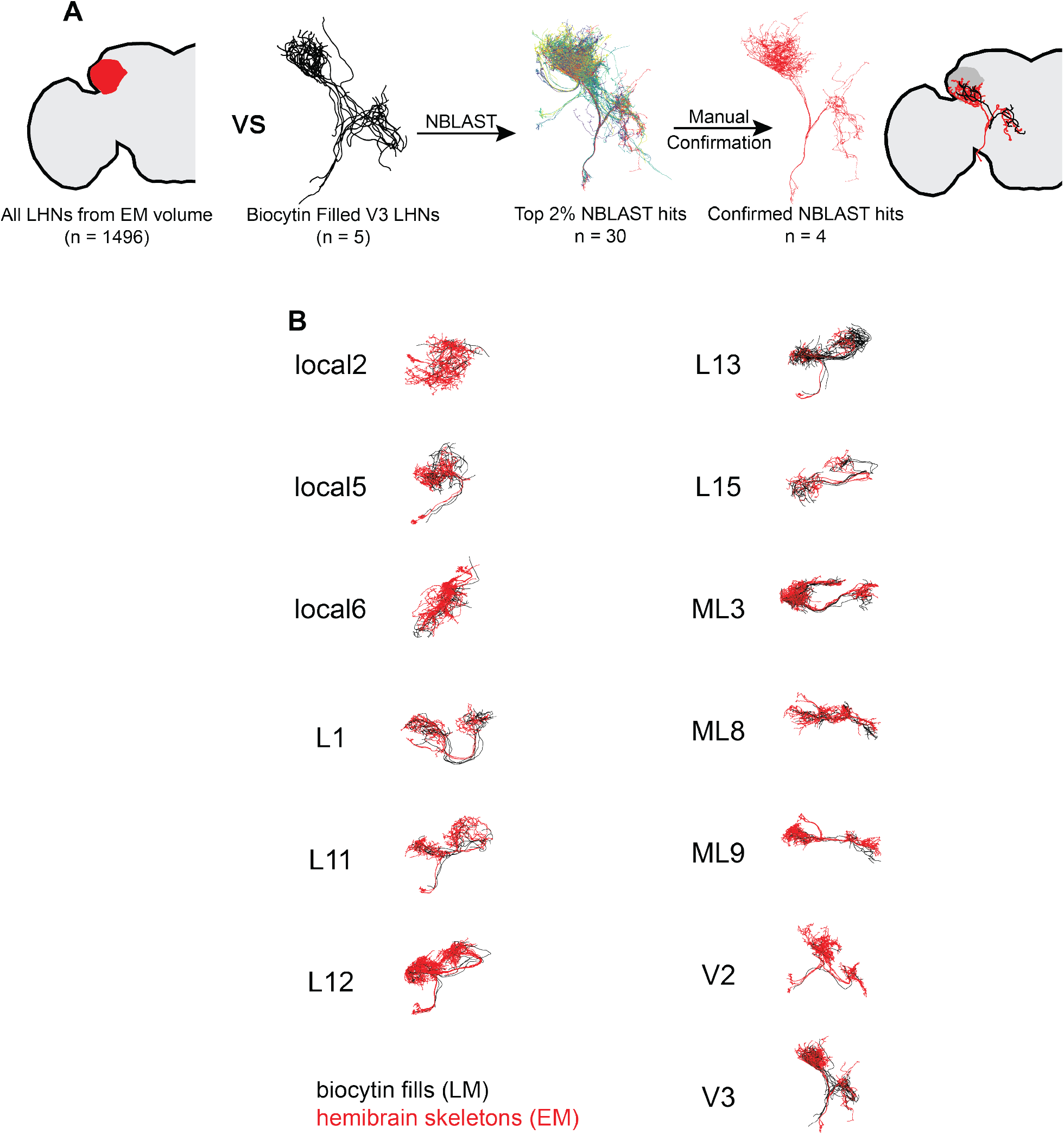
Identifying LHNs between EM and LM data (related to Figure 1) **(A)** Graphical illustration of procedure for finding the same LHN types between EM and LM datasets. We began with the skeletonized morphologies of all 1496 LHNs in the hemibrain connectome. For each LHN type as defined by Jeanne et al. (2018), we collected all skeletonized biocytin fills (the V3 LHN type is illustrated here) and quantitatively compared them to all hemibrain LHNs using the NBLAST algorithm. The highest scoring 30 hits from the hemibrain were selected for expert manual confirmation. **(B)** Superimposed representations of biocytin fills for each LHN type (black) and hemibrain skeletons (red).

**Figure S2:**
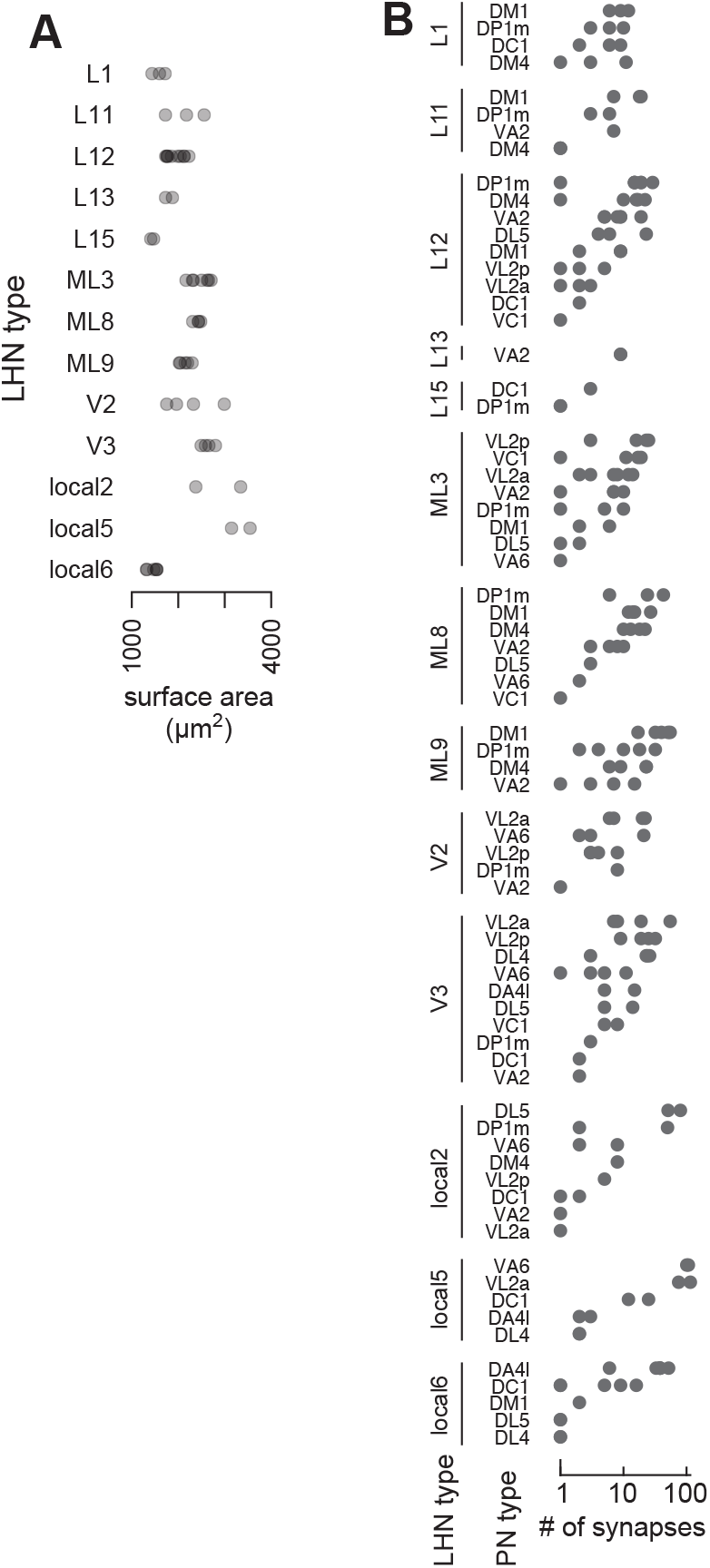
Characteristics of individual LHNs and PN-LHN connections (related to Figure 1) **(A)** LHN types have characteristic surface areas. Each point denotes a single neuron from the hemibrain connectome, grouped by LHN type. Variability between average surface areas across LHN types is greater than within types (ANOVA, F = 15.2, p = 1.8 × 10^−11^). **(B)** PN-LHN connections have characteristic synapse counts. Each point denotes a single PN-LHN connection, grouped by connection type. Variability between average synapse counts across types is greater than within types (ANOVA, F = 15.65, p = 2.6 × 10^−63^).

**Figure S3:**
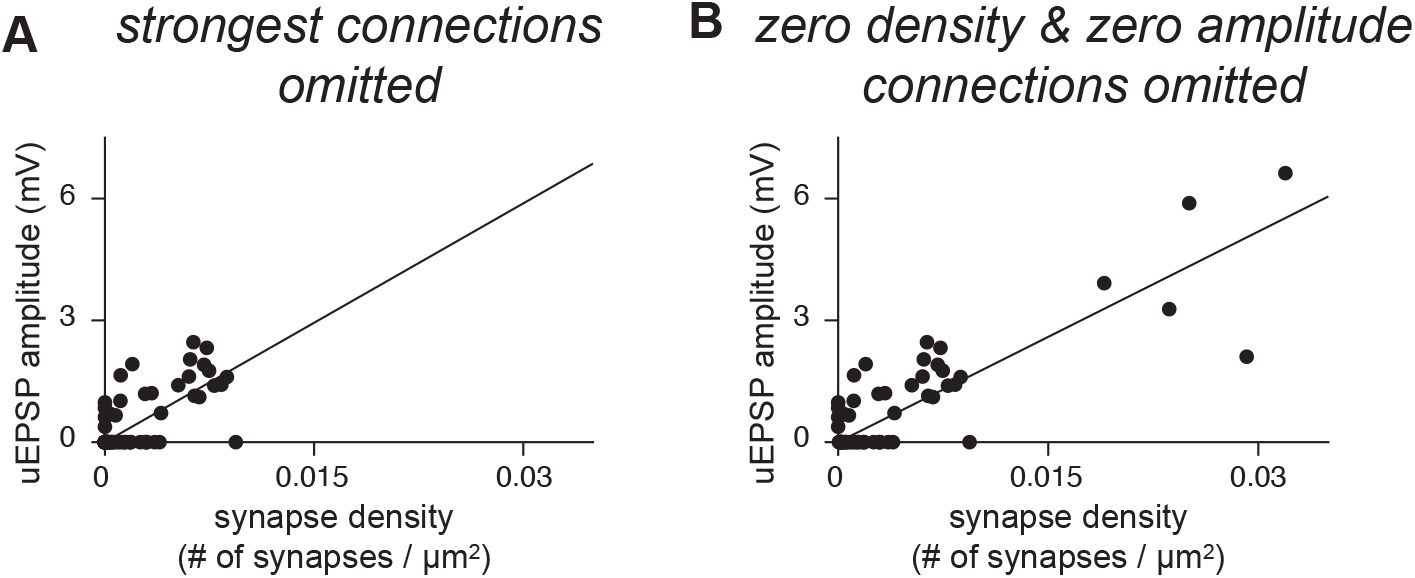
Further analysis of correspondence between anatomy and physiology of connections (related to Figure 3) **(A)** The correlation between synapse density and uEPSP amplitude persists in the absence of the 5 strongest connections (r^2^ = 0.55, p = 1.8 × 10^−24^). **(B)** The correlation between synapse density and uEPSP amplitude persists in the absence of the connections with zero density and zero amplitude (r^2^ = 0.72, p = 3.8 × 10^−18^). Note that a handful of PN-LHN pairs with 0 mV uEPSP amplitude have small (but non-zero) synapse densities, which are evident near the origin of this plot.

**Figure S4:**
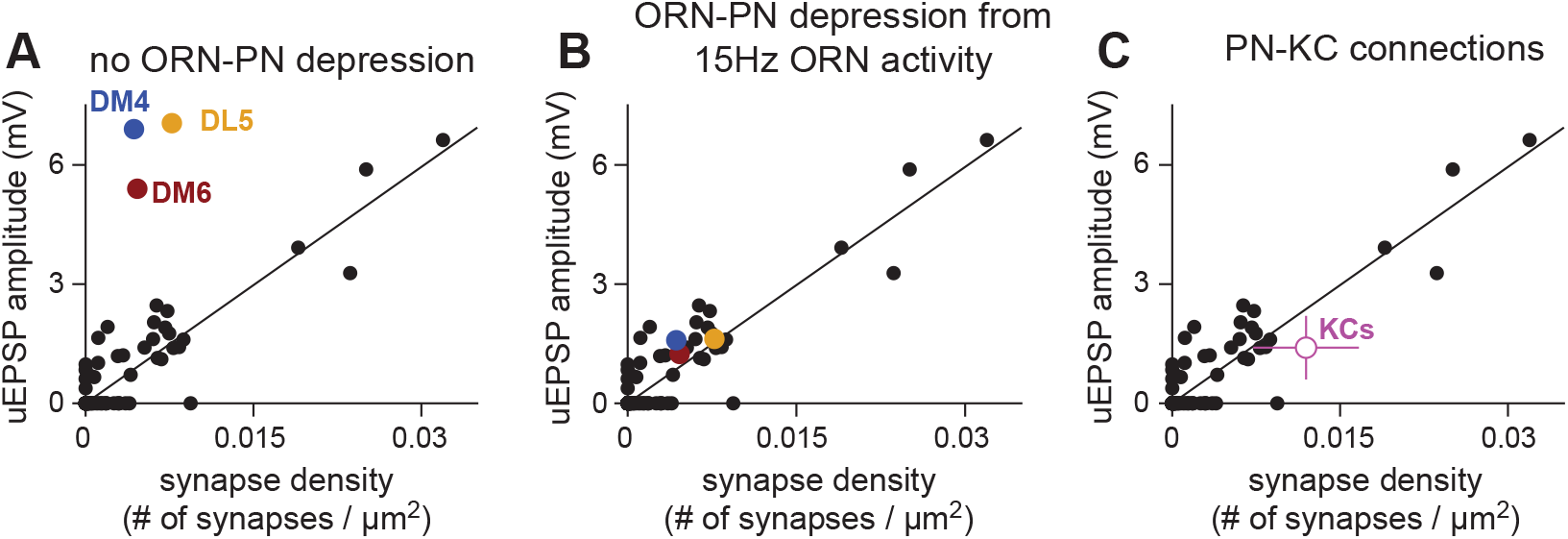
Relationship between synapse density and uEPSP amplitude is consistent with other cell classes (related to Figure 3) To determine whether the relationship we observe between synapse density and uEPSP amplitude for PN-LHN pairs is similar for other classes of neuron, we compared our data to those of two different classes of Drosophila synaptic connections: from olfactory receptor neurons (ORNs) to PNs and from PNs to Kenyon cells (KCs). **(A)** uEPSPs for three different ORN-PN connections were analyzed from previously published data (Kazama and Wilson, 2008) and compared to their corresponding synaptic density measurements, computed using identical methodology as for PN-LHN connections. A fourth ORN-PN connection (VM2) was not analyzed because that glomerulus was partially cut off in the hemibrain EM volume (Schlegel et al., 2021), precluding accurate estimates of surface area for the VM2 PN. The uEPSP amplitudes for each of these ORN-PN connections are much larger than for PN-LHN connections of the same synaptic density. However, ORN-PN synapses experience profound short-term depression and these uEPSPs were measured after long periods of ORN quiescence (via a nerve shock protocol without ORN spontaneous activity; Kazama and Wilson, 2008). **(B)** To estimate how activity would depress each of these uEPSP amplitudes, we implemented a simple model of short-term depression previously fit to ORN-PN synaptic connections (Nagel et al., 2015). Each ORN was simulated at a rate of 15Hz (the average rate of PN stimulation in Jeanne et al., 2018). At these rates, synaptic depression brings uEPSP amplitudes closer to those predicted from PN-LHN connections. **(C)** We compared PN-LHN connections to PN-KC connections. The mean and standard deviation of these uEPSP amplitudes were obtained from published measurements from whole-cell recordings of KCs (Turner et al., 2008). Synapse densities were obtained for all 10,739 connections from the 102 cholinergic uniglomerular PNs onto the 1927 traced KCs reported in the hemibrain connectome. Error bars denote standard deviation. uEPSP amplitudes for PN-KC connections were highly consistent with those for PN-LHN connections, although modestly lower than predicted. Overall, this comparison indicates that the relationship between synapse density and uEPSP amplitude is broadly similar across cell classes, but also highlights the importance of short-term synaptic plasticity in this relationship.

---

**Figure S5:**
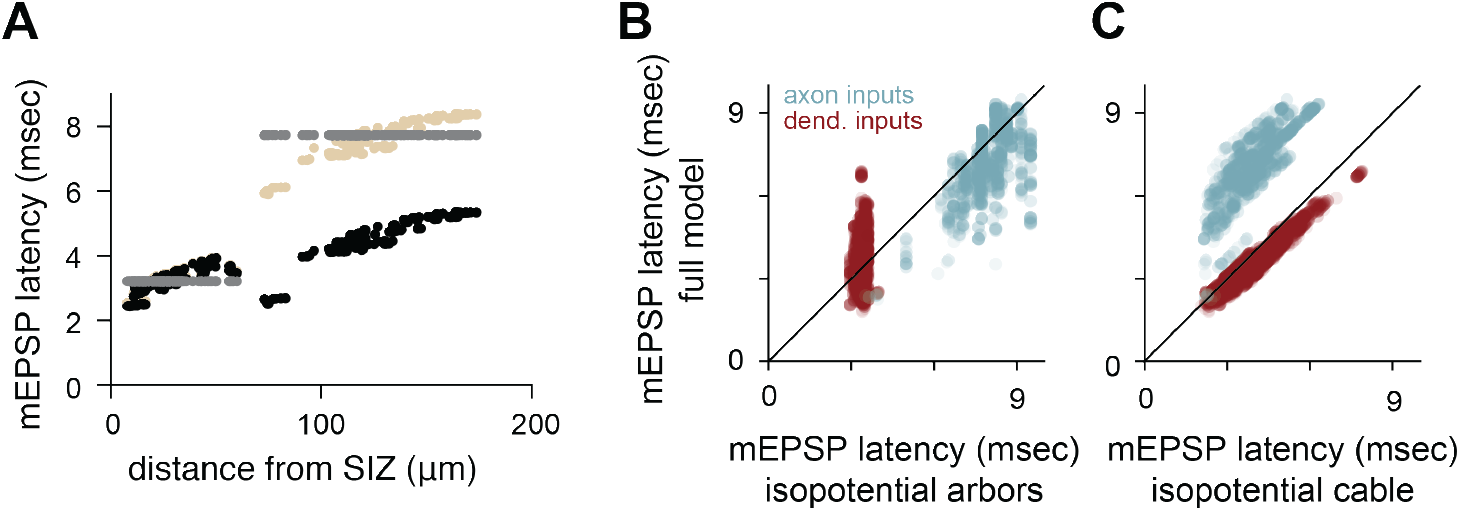
Democratization and division of mEPSP peak latencies (related to Figure 5) **(A)** Latencies to peak mEPSP for all ePN synapses onto the example local5 LHN, as a function of distance from the SIZ (tan). While the most proximal axonal synapses are nearly the same distance from the SIZ as the most distal dendritic synapses (∼60 μm), they evoke strikingly different mEPSP latencies. Grey points are the same measurements, but for a model with isopotential arbors. Black points are for a model with an isopotential inter-arbor cable. **(B)** Latencies to peak mEPSP simulated with the full model vs. the isopotential model for all synapses across 49 local LHNs. **(C)** Latencies to peak mEPSP simulated with the full model vs. the isopotential arbor model for all synapses across 49 local LHNs.

**Figure S6:**
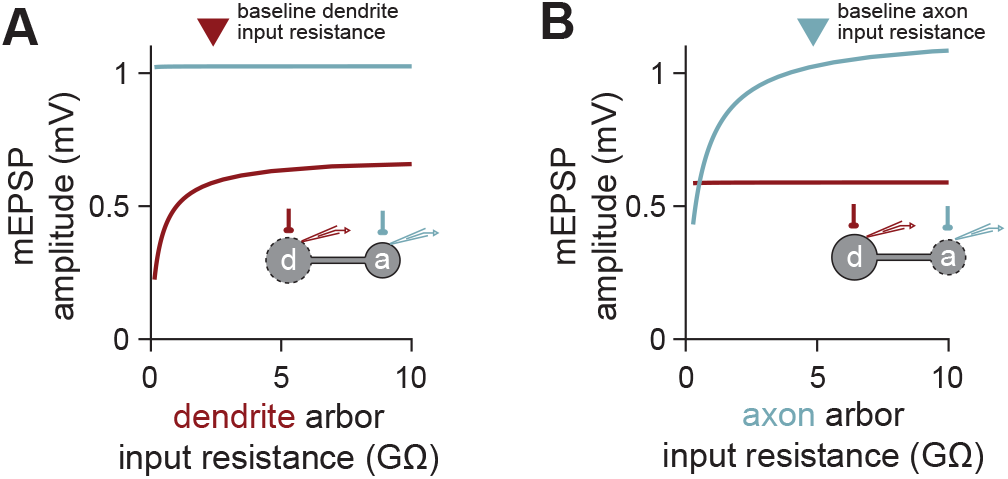
Input resistance of remote arbor does not impact input gain of local arbor (related to Figure 7) **(A)** Changing the dendrite arbor input resistance in the barbell model of an average local LHN does not impact input gain (mEPSP amplitude) in the axon arbor (light blue line), although it does change input gain within the dendrite arbor (red line). In each case, stimulation and measurement occur in the same arbor. **(B)** As in (A), but for changing the axon arbor input resistance.

**Figure S7:**
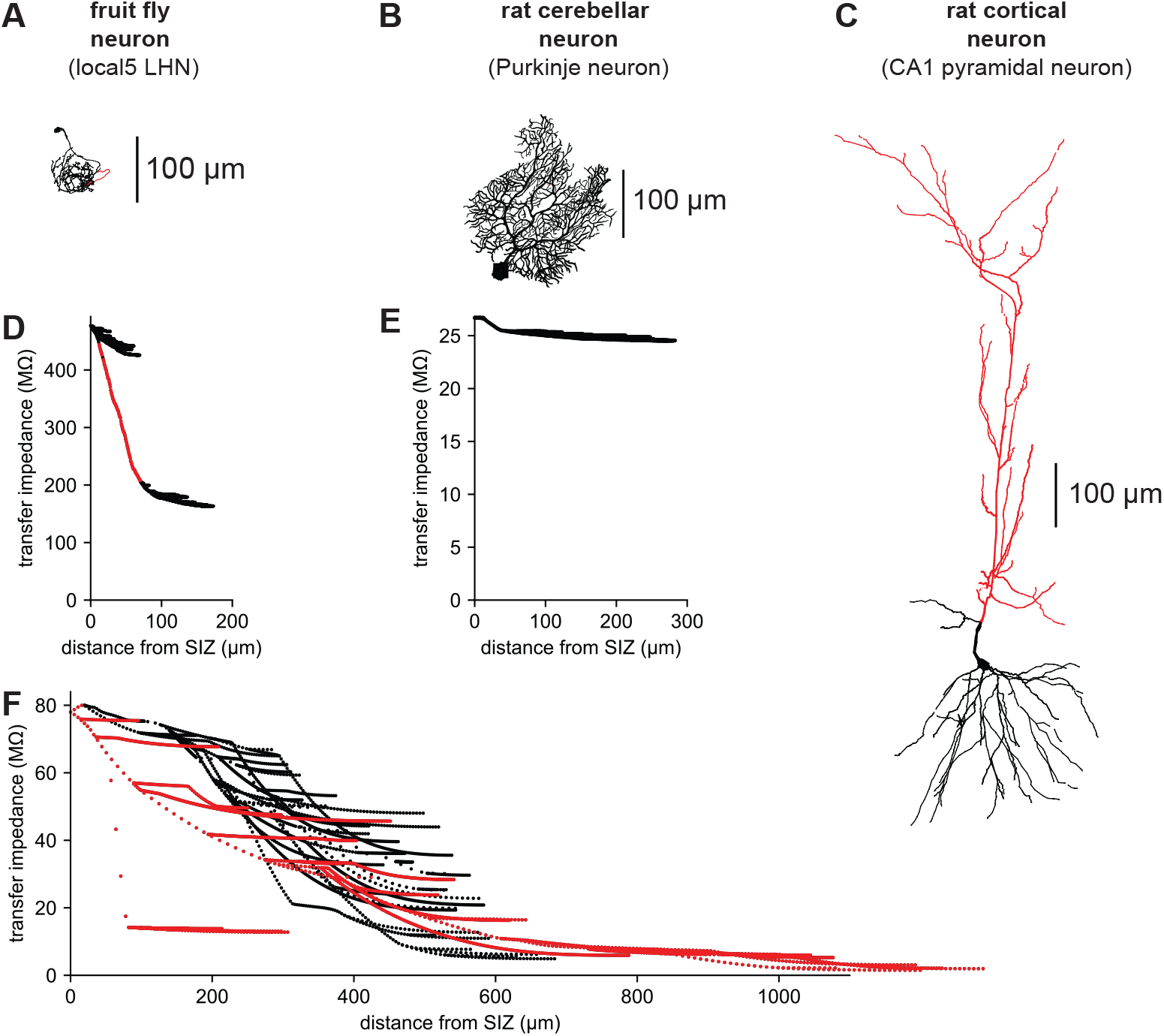
Comparison of transfer impedance profiles across species (related to Figure 8) Transfer impedance is a measure of synaptic efficacy, measured as the ratio of amplitude of voltage at the soma in response to an injection of current at another location. We calculated transfer impedance at 20 Hz, to provide a physiologically relevant measure of synaptic efficacy (Jaffe and Carnevale 1999). All plots are measured from the spike initiation zone, assumed to be at the soma for mammalian neurons. **(A-C)** morphologies of three distinct neuron types. **(D)** Transfer impedance profile of a local LHN, local5. The inter-arbor cable connecting the dendritic (higher transfer impedance) and axonal (lower transfer impedance) arbors is highlighted in red, corresponding to the red segment in (A). Transfer impedance drops rapidly along the inter-arbor cable, while it remains relatively stable within each arbor. **(E)** Transfer impedance profile of a rat Purkinje cell dendritic tree (Roth and Hausser, 2001). Transfer impedance is largely constant throughout the entire tree. **(F)** Transfer impedance profile of a rat CA1 pyramidal cell (Pyapali and Turner, 1996). The apical dendritic subtree is highlighted in red in both the morphological projection (top) and the transfer impedance graph (bottom). Transfer impedance is highly variable across dendritic locations in this cell type. The individual arbors of the fly neuron and the dendritic arbor of the Purkinje neuron exhibit minimal variability in transfer impedance, creating passive conditions for democratic integration of synaptic inputs. This arises, in part, because extensive branching creates many relatively short (sealed end) neurites. The high local input impedance of these neurites causes large local depolarizations that largely compensate for the additional cable filtering imposed by their length. In contrast, transfer impedance drops rapidly along the fly neuron inter-arbor cable and the rat cortical neuron apical dendritic trunk. These long cables (with open ends) exhibit fairly uniform local input resistance along their length, so they mostly cause passive attenuation and are well-approximated by an infinite cable. These minimally branched cables subdivide both fly and rat neurons into compartments with different synaptic efficacies. Biophysical parameters in the local5 LHN are as in the rest of this study. Parameters for the Purkinje neuron are based on Roth and Hausser (2001): r_a_ = 115 Ohm-cm, cm = 0.77 uF/cm2, g_L_ = 8.2 × 10-6 S/cm^2^. Parameters for the CA1 pyramidal neuron are based on Jaffe and Carnevale (1999): r_a_ = 200 Ohm-cm, cm = 1.0 uF/cm^2,^ g_L_ = 2.2 × 10-5 S/cm^2^.

